# Epigenetic and 3D genome reprogramming during the aging of human hippocampus

**DOI:** 10.1101/2024.10.14.618338

**Authors:** Nathan R. Zemke, Seoyeon Lee, Sainath Mamde, Bing Yang, Nicole Berchtold, B. Maximiliano Garduño, Hannah S. Indralingam, Weronika M. Bartosik, Pik Ki Lau, Keyi Dong, Amanda Yang, Yasmine Tani, Chumo Chen, Qiurui Zeng, Varun Ajith, Liqi Tong, Chanrung Seng, Daofeng Li, Ting Wang, Xiangmin Xu, Bing Ren

## Abstract

Age-related cognitive decline is associated with altered physiology of the hippocampus. While changes in gene expression have been observed in aging brain, the regulatory mechanisms underlying these changes remain underexplored. We generated single-nucleus gene expression, chromatin accessibility, DNA methylation, and 3D genome data from 40 human hippocampal tissues spanning adult lifespan. We observed a striking loss of astrocytes, OPC, and endothelial cells during aging, including astrocytes that play a role in regulating synapses. Microglia undergo a dramatic switch from a homeostatic state to a primed inflammatory state through DNA methylome and 3D genome reprogramming. Aged cells experience erosion of their 3D genome architecture. Our study identifies age-associated changes in cell types/states and gene regulatory features that provide insight into cognitive decline during human aging.

## Introduction

Age is the greatest risk factor for neurodegenerative diseases such as Alzheimer’s disease, Parkinson’s disease, Huntington’s disease, and amyotrophic lateral sclerosis (*1*). As 23% of the US population will be older than 65 by the year 2054 (*2*), there is an increasing necessity to understand how the normal aging process promotes disease. Every multicellular organism undergoes the aging process, leading to gradual changes in many biological systems and diminishing tissue and cellular homeostasis. As the human brain ages, it presents physiological changes that are associated with a decline in cognitive function (*3, 4*) which can greatly diminish quality of life. The hippocampus is a brain region of particular interest for understanding age-related cognitive dysfunction, due to its role in learning, episodic memory, emotional regulation, and spatial navigation—all behaviors affected in normal aging (*5*). In addition, hippocampal atrophy occurs during normal aging and is associated with memory impairment (*3, 4*).

At the cellular and molecular levels, hallmarks of aging include changes in intercellular communication, inflammation, mitochondrial dysfunction, and epigenetic changes such as DNA methylation and chromatin remodeling (*6*). These alterations contribute to age-related impaired regulation of gene expression and other cellular processes (*6*). However, when these changes occur and how they are regulated during aging remain unanswered. Since cellular phenotypes of aging are cell type-specific, it is imperative we generate molecular profiles of each cell type in order to accurately characterize age-related gene expression alterations and epigenetic changes. Recent research has uncovered cell type-specific changes with aging in human brain, including alterations in gene expression and chromatin accessibility in prefrontal cortex (*7*), gene expression and DNA methylation in cortical neurons (*8*), and changes to the 3D genome structure of cerebellar cells (*9*). However, despite its crucial role in cognitive function, a detailed investigation of single-cell epigenomics and 3D genome architecture in the aging human hippocampus is still lacking. In this study, we provide a comprehensive analysis of cell type-specific changes in the aging human hippocampus by profiling 3D genome organization alongside DNA methylation, gene expression, and chromatin accessibility. The analysis of these four molecular modalities from the same cell types revealed age-associated changes that were regulated in a complex yet coordinated manner across different layers of epigenetic mechanisms, as well as changes only captured by individual modalities.

Specifically, our study revealed age-related decline in astrocytes, including those involved in regulating synaptic connectivity. Strikingly, hippocampal microglia undergo an age-associated switch to an epigenetically encoded primed immune state through DNA methylome reprogramming. In addition, we discovered age-associated reorganization of 3D genome architecture that coincided with changes in gene expression and the chromatin accessible landscape. Our integrative multi-omic analysis provides novel molecular insights into cell type-specific aging phenotypes in the human brain with implications for how altered gene regulatory programs promote age-related cognitive dysfunction.

## Results

### Molecular and Cellular Dynamics of Human Hippocampus during Aging

We carried out our study using hippocampal tissues from 40 postmortem neurotypical human donors (Fig. 1A and table S1). To achieve an age and sex balanced cohort spanning the adult lifespan, we included 10 donors, 5 males and 5 females, from each of 4 age groups: 20-40, 40-60, 60-80, and 80-100 (Fig. 1A and table S1). To identify age-associated dynamics in the transcriptome, epigenome, and 3D genome organization of each cell type, we utilized two single-nucleus multi-omic sequencing assays: 10x multiome (10x Genomics) (Fig. 1B and fig. S1A) and single-nucleus methyl-3C sequencing, (snm3C-seq) (*10*), aka Methyl-HiC (*11*), (Fig. 1C and fig. S1B). From each nucleus, 10x multiome generates both snRNA-seq and snATAC-seq data, while snm3C-seq generates both DNA methylome and chromatin contact data.

**Fig. 1.**
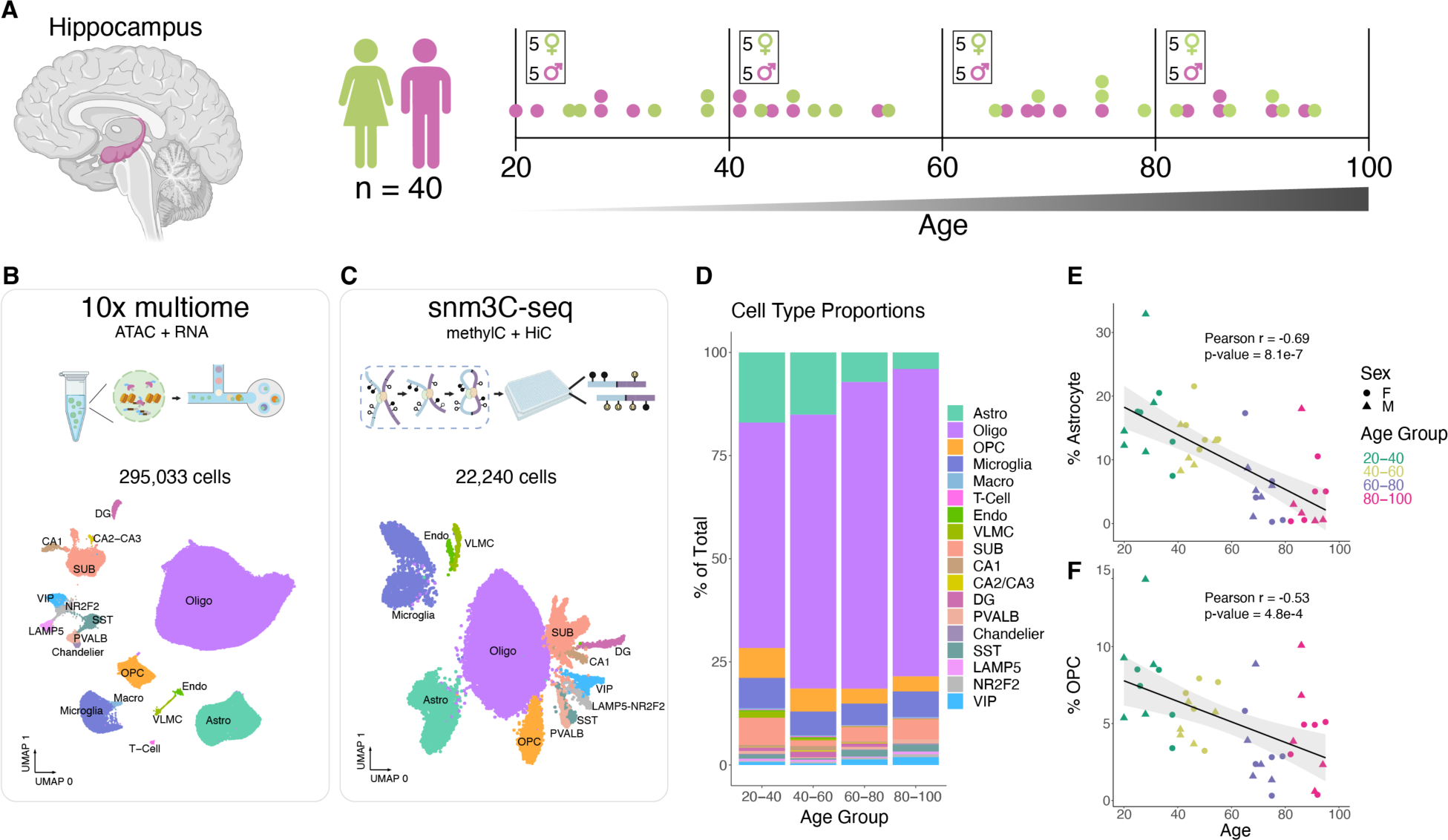
Cellular composition of the aging human hippocampus. (**A**) Diagram of the human hippocampus, generated with BioRender. (**B** and **C**) Workflow diagrams (made with BioRender) and uniform manifold approximation and projection (UMAP) embeddings of 10x multiome RNA-seq (B) and snm3C-seq DNA methylation (C) clustering, colored and annotated by cell type. OPC, oligodendrocyte precursor cells; VLMC, vascular leptomeningeal cells. (**D**) Stacked bar plots representing percentage of cell type recovered from donors in indicated age group. (**E** and **F**) Scatter plots showing the percentage of recovered astrocytes (E) or OPC (F) out of total cells vs donor age from 10x multiome experiments.

After filtering out low quality nuclei (methods), we retained 295,033 nuclei with 10x multiome and performed in-depth analysis of their transcriptomes and chromatin accessibility (Fig. 1B and fig. S1, A,C, and D). Using known marker gene expression (fig. S2A) and reference mapping to a published snRNA-seq dataset (*12*), we annotated clusters corresponding to 18 subclasses of cells (Fig. 1B). Similarly, we mapped DNA methylomes and 3D genome folding from 22,240 nuclei (fig. S1, B, E and F), and grouped them into 13 subclasses (Fig. 1C) using hypomethylation levels of known marker genes and integration with our annotated snRNA-seq dataset (fig. S2, B to D). Together, this provided us cell type-resolved transcriptomes, chromatin accessible profiles, DNA methylomes, and 3D genome profiles in hippocampal cells from 40 donors (fig. S2, E to K), with 39 out of 40 donors matching between 10x multiome and snm3C-seq (table S1). These data are available for visualization as a data hub on the WashU Epigenome Browser: https://epigenome.wustl.edu/seahorse/.

Neurons generally had more transcripts detected, genes expressed, and chromatin accessible DNA fragments captured compared to non-neurons (fig. S2, E to G). Consistent with other reports (*13, 14*), we observed higher levels of non-CG methylated sites (mCH) in neuronal cell types than in glial cells (fig. S2I).

We first looked for cell types that change in abundance during aging (Fig. 1D, fig. S3, and table S2). In particular, hippocampal astrocytes displayed the most age-correlated decline (PCC = -0.68, p-value 1.1e-6) (Fig. 1E and fig. S3, A and B). Additionally, both oligodendrocyte precursor cells (OPC) and endothelial cell abundance significantly decreased as donor age increased (Fig. 1F and fig. S3C). The age-correlated decrease in OPC is consistent with a study of the human cortex (*7*). While the age-correlated decrease in endothelial cells is significant (PCC = -0.52, p-value = 4.8e-4), we note that the number of recovered cells per donor is very low (< 1%) and highly variable (fig. S3A). Nonetheless, loss of hippocampal endothelial cells and astrocytes with age could contribute to the documented breakdown of blood-brain barrier with age (*15*). Similar age-correlated decline in the number of astrocytes, OPC, and endothelial cells was observed in our snm3C-seq data (fig. S3, D to G, and table S3).

The observed decline in the number of astrocytes with age was particularly intriguing considering their indispensable roles in neuronal signaling in the adult brain (*16*). We confirmed this decline using confocal imaging of human hippocampal CA3 tissue by immuno-staining for astrocytic marker ALDH1L1 (Fig. 2, A and B).

**Fig. 2.**
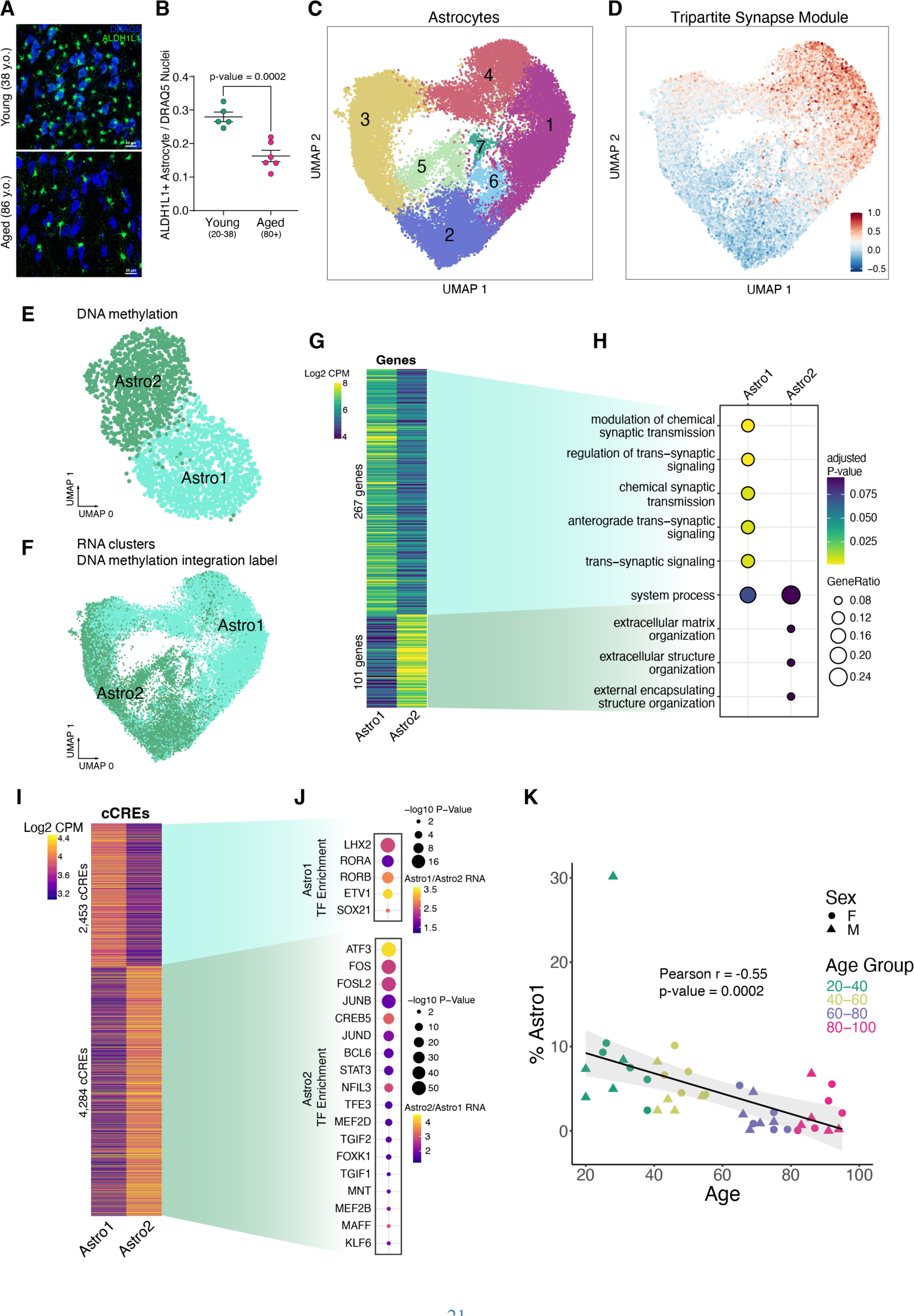
Hippocampal tripartite astrocytes. (**A**) Confocal imaging of hippocampal CA3 tissue immuno-stained for the astrocytic marker ALDH1L1 (green) and the nuclei marker DRAQ5 (blue). (**B**) Ratio of ALDH1L1-positive astrocytes to DRAQ5-stained nuclei in young donors (n=5, 20-38 years old) and aged donors (n=6, 80+ years old). P-value = 0.0002 from an unpaired Welch’s T-test. (**C**) UMAP plot of astrocytes based on 10x multiome RNA data, annotated with 7 astrocyte subclusters. (**D**) UMAP showing the average module score (methods) of 18 tripartite synaptic genes (*19*). (**E**) UMAP plot of astrocytes from snm3C-seq DNA methylation data, annotated with 2 astrocyte subclusters. (**F**) UMAP plot of astrocytes from 10x multiome RNA data, annotated with 2 DNA methylation astrocyte subclusters by label transfer using hypomethylation gene scores and RNA integration. (**G**) Heatmap displaying Log2 RNA CPM of Astro1 and Astro2 marker genes. (**H**) Gene ontology terms for enriched biological processes of Astro1 and Astro2 marker genes. (**I**) Heatmap displaying Log2 ATAC-seq fragments CPM of Astro1 and Astro2 marker cCREs. (**J**) Enriched TF motifs for Astro1 and Astro2 marker cCREs that also exhibited up-regulated TF expression in the indicated subcluster. (**K**) Scatter plots showing the percentage of recovered Astro1 cells out of total cells vs. donor age from 10x multiome experiments.

Given their known functional diversity, we sub-clustered astrocytes to resolve different functional cell states (Fig. 2C). For example, a subset of astrocytes function in synaptic maintenance by directly clearing neurotransmitters in the synaptic cleft and signaling to proximal neurons via gliotransmitters at tripartite synapses (*17, 18*). Tripartite astrocytes express gene markers that can be used to distinguish them from non-tripartite astrocytes (*19*). Two astrocyte subclusters, 1 and 4 (Fig. 2C), were enriched for expression of a set of 18 tripartite astrocyte genes (*19*) (Fig. 2D, Methods). We also subclustered astrocytes using their DNA methylation profiles, and identified 2 subclusters, annotated as “Astro1” and “Astro2” (Fig. 2E). Cells in the Astro1 subcluster were more hypomethylated at CG dinucleotides within the gene bodies of tripartite synapse astrocyte genes, suggesting higher expression of these genes compared to cells in the “Astro2” subcluster (fig. S4A). We integrated DNA methylation with RNA using hypomethylation gene scores (methods), and found the Astro1 DNA methylation cluster aligned with RNA clusters 1 and 4 (Fig. 2F), providing further evidence that the Astro1 subcluster represents tripartite astrocytes. For consistency across modalities, we labeled tripartite 10x multiome astrocyte clusters 1 and 4 as “Astro1” and non-tripartite astrocyte clusters 2, 3, 5, 6, and 7 as “Astro2” (Fig. 2C).

We identified genes differentially expressed between Astro1 and Astro2 (Fig. 2G and table S4), and observed gene ontology enriched terms related to synaptic signaling for Astro1, and gene ontology enriched terms related to extracellular matrix organization enriched for Astro2 (Fig. 2H). We also identified 6,737 candidate cis-regulatory elements (cCREs) that were significantly more accessible in Astro1 than in Astro2, or vice versa (Fig. 2I and table S5). The Astro1 cCREs were enriched for TF motifs corresponding to LHX2, RORA, RORB, ETV1, and SOX21 binding, suggesting a potential role for these TFs in promoting tripartite astrocyte cell state (Fig. 2J). As with total astrocytes, the synaptic-signaling tripartite astrocytes exhibited an age-correlated drop in their number with age (Fig. 2K).

### Age-correlated Changes in Epigenome and Gene Regulatory Programs in Hippocampal Astrocytes

We utilized our age-balanced cohort to identify genes whose expression significantly correlated with age in hippocampal astrocytes and other cell types. For every cell type, we calculated Pearson correlation coefficients (PCC) between expression levels of each gene with donor age (Fig. 3A) and identified genes that are either negatively or positively age-correlated (Fig. 3B, fig. S5A, and table S6). We found the negatively age-correlated genes in astrocytes were highly enriched for genes that function in the mitochondria for ATP synthesis (Fig. 3C). These functions were also enriched in genes negatively age-correlated in subiculum (SUB) excitatory neurons as well as Chandelier, LAMP5, and VIP inhibitory neurons (fig. S5B). As part of mitochondrial dysfunction surveillance, cells respond to depleted ATP by activating lysosomal autophagy (*20, 21*). Consistent with this interplay, genes with increased transcription in aged astrocytes were enriched for lysosomal microautophagy (Fig. 3D and fig. S5C), potentially leading to “autophagy-dependent cell death” (*22*) in aging hippocampal astrocytes. Consistent with a recent study of prefrontal cortical astrocytes (*23*), genes involved in the synaptic neuron and astrocyte program (SNAP) showed decreased expression in hippocampal astrocytes during aging (fig. S4B).

**Fig. 3.**
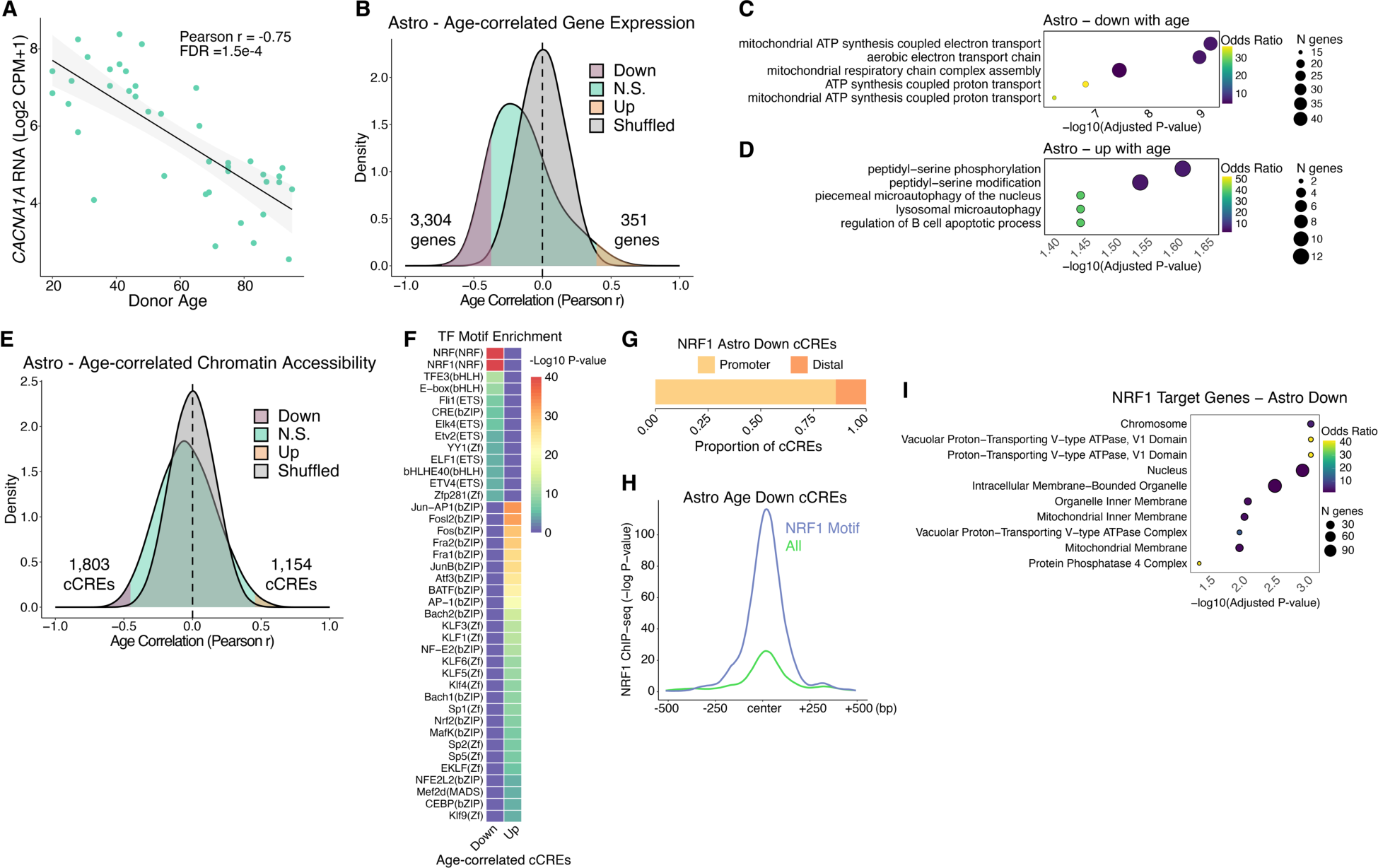
Aging gene regulatory programs in astrocytes. (**A**) Scatter plot of *ZNF804A* expression vs. donor age. (**B**) Density plots for Pearson r values of age vs. expression (log2 CPM+1) across 40 donors. Shuffled represents a null distribution of shuffling the expression values of the donors for each gene. (**C** and **D**) Gene ontology dot plots displaying significant biological process enriched terms for age-correlated genes that decrease with age (C) or increase with age (D) in astrocytes. (**E**) Density plots for Pearson r values of age vs. chromatin accessibility (log2 CPM+1) across 40 donors. (**F**) TF motif enrichment heatmap for significant enrichments, q-value < 0.05. “Up” indicates chromatin accessibility increases with age and “Down” indicates chromatin accessibility decreases with age. (**G**) Ratio of cCREs with negatively age-correlated accessibility in astrocytes with an NRF1 TF motif that are either promoter proximal (<1 kb) or promoter distal (> 1kb) from a protein-coding transcription start site. (**H**) Average -Log p-value NRF1 ChIP-seq signal in SK-N-SH cell line (ENCODE Accession ENCFF762LEE) at all astrocyte cCREs that decrease accessibility with age (n = 1803, yellow line) or the subset that contain an NRF1 TF motif (n = 354, blue line). (**I**) Gene ontology biological process significantly enriched terms (q-value < 0.05) for genes that have negative age-correlated accessibility in astrocytes with an NRF1 TF motif at their promoter.

To identify the transcriptional regulators involved in the above age-associated gene expression programs in astrocytes, we examined cCREs that exhibit age-correlated chromatin accessibility in astrocytes (Fig. 3E, fig. S6A, and tables S7 and S8). These resided mostly at promoter distal genomic regions (fig. S6B). To find putative TFs with age-correlated DNA binding activity, we performed TF motif enrichment on age-correlated cCREs (Fig. 3F, fig. S6C, and table S9). We found bZIP family TF motifs enriched in astrocyte cCREs gaining accessibility with age, such as AP-1/ATF superfamily members (Fig. 3F and fig. S6C). At cCREs that lose accessibility in aging astrocytes, we observed highest enrichment for nuclear respiratory factor (NRF) TF motifs (Fig. 3F and fig. S6C). This included binding sites for nuclear respiratory factor-1 (NRF1), which is known to activate nuclear genes with mitochondrial functions (*24*). Given that we identified ∼40 genes having mitochondrial function, such as ATP synthesis, that lose expression with age in astrocytes (Fig. 3C), we hypothesized that reduced DNA binding of NRF1 may be responsible. Consistent with this hypothesis, we found that cCREs with age-reduced accessibility in astrocytes containing an NRF1 motif (table S10) are enriched at promoters (Fig. 3G), many of which were previously shown to bind NRF1 in human neuroblastoma cells (*25*) (Fig. 3H). Additionally, the genes with both reduced accessibility with age in astrocytes and NRF1 motifs in their promoters (table S11) encode ATPases and mitochondrial membrane proteins (Fig. 3I). Taken together, these lines of evidence suggest a potential mechanism where reduced NRF1 DNA binding leads to mitochondrial dysfunction, reduced ATP levels and increased likelihood of autophagy-dependent cell death in aging human hippocampal astrocytes. However, NRF1 mRNA levels were unaltered during aging (PCC = -0.05, p-value = 0.76), suggesting its reduced DNA binding is regulated at the protein level.

### Proinflammatory Gene Regulatory Programs in Microglia During Aging

Compared to other cell types, microglia possessed the most genes with positive age-correlated expression (Fig. 4A and fig. S5A). These genes were enriched for immune response terms such as regulation of phagocytosis, response to cytokine, and response to interferon-gamma (Fig. 4B), and likely contribute to neuroinflammation of the aging hippocampus (*26*). Microglia also had a relatively large number of age-correlated cCREs (Fig. 4C and fig. S6A), demonstrating that their epigenomes are highly dynamic during aging. While specific TF motifs were not significantly enriched in the cCREs that lose accessibility with age, we found several TF motifs enriched at cCREs that gain accessibility with age (Fig. 4D and fig. S6C). Enrichments were highest for bZIP and ETS family members and included TFs that mediate proinflammatory responses in microglia such as CEBP, AP-1, STAT family, and p53 (Fig. 4D and fig. S6C) (*27, 28*).

**Fig 4.**
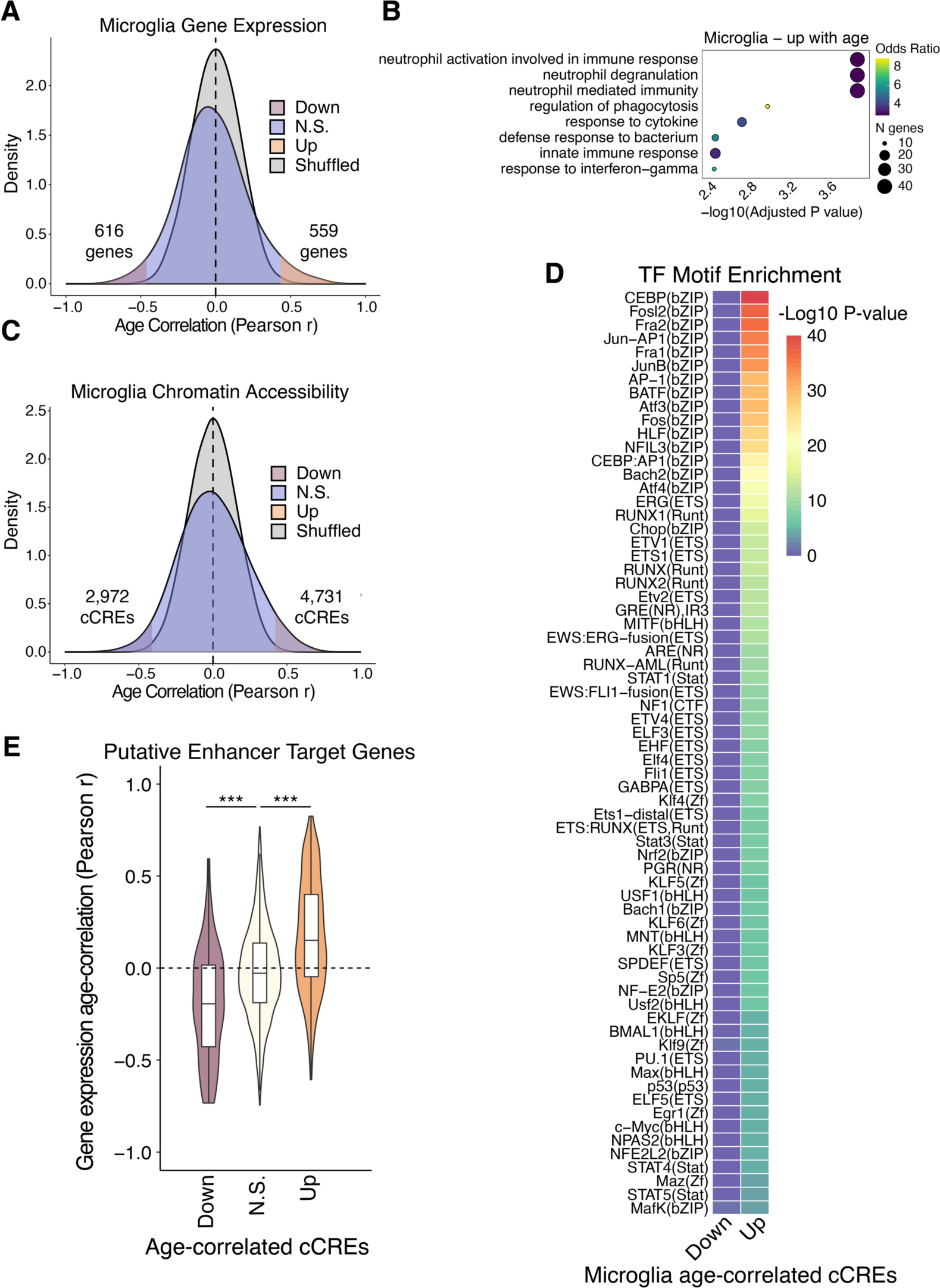
Aging gene regulatory programs in microglia. (**A**) Density plots for Pearson r values of age vs. expression (log2 CPM+1) across 40 donors. Shuffled represents a null distribution of shuffling the expression values of the donors for each gene. (**B**) Gene ontology dot plots displaying significant biological process enriched terms for age-correlated genes that increase with age in microglia. (**C**) Density plots for Pearson r values of age vs. chromatin accessibility (log2 CPM+1) across 40 donors. (**D**) TF motif enrichment heatmap for significant enrichments, q-value < 0.05. “Up” indicates chromatin accessibility increases with age and “Down” indicates chromatin accessibility decreases with age. (**E**) Violin plots showing the distribution of Pearson correlation coefficients for age-correlation of expression for target genes of predicted ABC enhancers. P-values from two-sided, unpaired Wilcoxon-rank sum test. * p < 0.05, ** p < 0.01, *** p < 0.001

In order to investigate the potential influence of age-correlated cCREs on the aging transcriptome, we predicted enhancers and their putative target genes from chromatin accessibility and Hi-C data of each hippocampal cell type using the activity-by-contact (ABC) model (*29*). This yielded 36,275 promoter distal (> 1 kilobase) putative enhancers (table S12) targeting 14,135 unique genes that displayed matching cell type-specific activities across hippocampal cell types (fig. S6, D and E). We observed significant correspondence between putative enhancers that have age-correlated chromatin accessibility with the age-correlation of their target gene’s expression (Fig. 4E and fig. S6F). This correspondence suggests that age-related changes to the epigenome have a major influence on the age-related transcriptome.

### Epigenetic Priming of Aging Microglia through a DNA Methylome Reprogramming Event

Given the high degree of age-dynamic chromatin accessibility in microglia, we explored the consequence of aging on microglial DNA methylomes using methylated fractions at all sequenced CG dinucleotides (mCG). We identified two robust microglia subclusters labeled “Micro1” and “Micro2”, that were consistently detected with or without batch correction (Fig. 5A and fig. S7A). We found a striking age-dependence of the proportion of Micro1 and Micro2 cells (Fig. 5B). 20-40 year old donors contributed mostly to Micro1, 80-100 year old donors contributed almost exclusively to Micro2, and the 40-60 and 60-80 age groups were more split between Micro1 and Micro2 in an age-dependent manner (Fig. 5B).

**Fig. 5.**
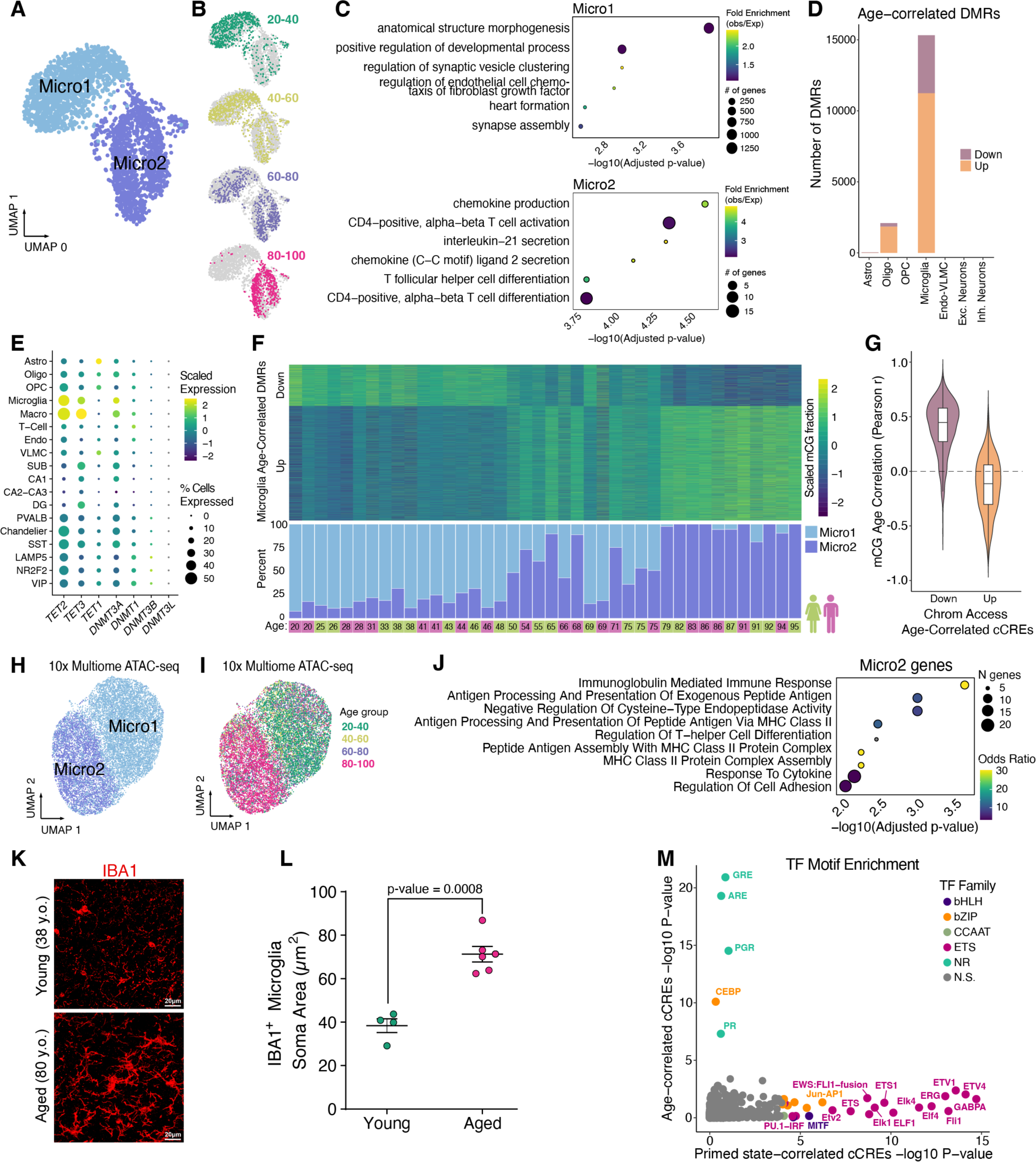
Epigenetic cell state switch in aging microglia. (**A**) UMAP plot of microglia from snm3C-seq, annotated with 2 DNA methylation microglia subclusters. (**B**) UMAP embeddings showing the distribution of microglia cells across different age groups. (**C**) Dot plots displaying gene ontology terms for enriched biological processes for the nearest genes (closest transcription start site) from DMRs hypomethylated in Micro1 (top) and Micro2 (bottom). (**D**) Stacked bar plots showing number of significant (FDR < 0.1) age-correlated DMRs in each subclass. (**E**) Dot plot showing the scaled average single cell gene expression levels of enzymes involved in DNA (de)methylation across cell types. The color of each dot represents the expression level and the size indicates the percentage of cells expressing the gene. (**F**) Heatmap of Age-correlated DMRs (rows), displaying row-scaled methylated cytosine proportions for each donor (columns) (top) and stacked bar plots showing the ratio of Micro1 to Micro2 cells recovered for each donor in snm3C-seq experiments (bottom). (**G**) Violin plots showing the distribution of Pearson correlation coefficients for age-correlation of mCG fractions at DMRs overlapping age-correlated cCREs. (**H**) UMAP displaying microglia 10x multiome clustered by ATAC-seq cCREs identified in microglia that overlap Micro1 or Micro2 DMRs (n = 3,405). Cells colored by Micro1 or Micro2 annotation. (**I**) Same as (H) except cells colored by age group. (**J**) Significant GO Biological Process enriched terms for Micro2 differentially upregulated genes compared to Micro1. (**K**) Representative micrographs of IBA immuno-stained hippocampus CA3 region in two human donors. (**L**) Dot plot showing quantification of microglia soma area between young (n=3; 20-38 y.o.) and aged (n=6; 80+ y.o.). P-value from linear mixed-effect model (LME). (**M**) Scatter plot displaying TF motif enrichment for cCREs either Micro2 percentage correlated (“Primed-state correlated cCREs”, fig. S13B) or age-correlated (fig. S13C) in 16 donors. Only TF motifs with q-value < 0.05 are colored by their TF family.

To further characterize these age-associated microglia subclusters, we determined subcluster-specific differentially methylated regions (DMRs) (table S13) with differential hypomethylation in either Micro1 or Micro2 (table S14). Micro1-hypomethylated DMRs were enriched with the binding motif of ZNF281 (Zfp281 in mouse) (fig. S7B), a transcription factor that regulates dynamic DNA methylation in early mammalian development (*30*). These DMRs are proximal to genes with developmental functions (Fig. 5C, top); many not expressed in the adult hippocampus yet hypermethylated during aging. For example, Homeobox A (HOXA) cluster genes had a strong correlation between DNA methylation and donor age (PCC = 0.84, p-value = 9.12e-12), which was unique to the microglia cell type (fig. S8, A to C). In contrast, Micro2-hypomethylated DMRs were enriched near genes involved in activated microglia functions such as chemokine production (Fig. 5C, bottom), and contained TF motifs of the bZIP and ETS families such as AP-1 and FLI1 (fig. S7B).

As with chromatin accessibility, microglia had relatively high numbers of age-correlated DMRs across cell types (Fig. 5D and table S15). Of these, 73% (11,239/15,327) became hypermethylated with age. Additionally, microglia showed strong correlation between their DNA methylation clock (*31*) predicted biological age and chronological age, while OPC and inhibitory neurons showed very low correlation (fig. S9A). Transcription levels of genes coding for enzymes directly involved in adding and removing DNA methylation, *DNMT3A, TET1*, and *TET2*, were highest in hippocampal microglia and macrophages (Fig. 5E), which may contribute to the high degree of dynamic CG methylation in these cell types during aging. The DNA methylation levels of age-correlated DMRs matched the ratio of Micro1 and Micro2 cells for each donor (Fig. 5F, fig. S8C, table S16), demonstrating that most age-correlated DNA methylation is a result of epigenetic cell state switching.

We observed a general correspondence between DNA methylation and chromatin accessibility at age-correlated cCREs in microglia (Fig. 5G and fig. S9B). Given this correspondence, we reasoned that chromatin accessible cCREs that overlap Micro1 and Micro2 DMRs (n = 3,405) may resolve Micro1 and Micro2 clusters in the 10x multiome microglia, where as gene expression and total cCREs failed to resolve these states (fig. S10). Indeed, when using these cCREs as features, two distinct, age-dependent clusters were apparent (Fig. 5, H and I). The proportion of cells between these clusters closely matched proportions obtained from snm3C-seq microglia cells across donors (PCC = 0.88, p-value = 1.9e-13) (fig. S11A).

We further characterized Micro1 and Micro2 by identifying genes that were differentially expressed between the cells in these two clusters using donor age as a latent variable (table S17). Genes that were upregulated in Micro2 were enriched for major histocompatibility complex class II (MHC II) encoded by the human leukocyte antigen genes (HLAs) (Fig. 5J and fig. S11, B and C). Elevated MHC II is a marker for primed microglia (*32–36*), known to reside in the hippocampus of aged rodents (*26, 32*). Upon various immune challenges, primed microglia have more pronounced proinflammatory responses than non-primed microglia including a transient, high induction of cytokines, sufficient for memory impairment (*26, 37, 38*). The upregulation of MHC II suggests that Micro2 cells represent primed microglia, established through a DNA methylation reprogramming event. While primed microglia are characterized by their exaggerated immune responses, how they retain their immune memory is unknown (*39*). Our data suggest that an epigenetic memory is stably encoded through reprogramming of the DNA methylome, producing cells that are primed to revert to the proinflammatory, activated cell state where high levels of cytokines are produced.

In addition to MHC II upregulation, primed microglia adopt a more dystrophic morphology with a larger soma (*36, 40*). We observed a dramatic difference in the morphology of microglia in the hippocampus of aged donors compared to young donors (Fig. 5K). The aged hippocampal microglia had morphological characteristics of primed microglia, such as a significantly larger soma (Fig. 5L and fig. S12). The morphological characteristics of the aged microglia, which are mostly Micro2, provides further evidence that Micro2 are primed microglia.

To determine putative drivers of microglia priming, we identified differentially accessible cCREs between Micro1 and Micro2 (table S18), and found them to be enriched for ETS-domain transcription factor binding motifs (fig. S11D). To further separate epigenomic changes during the switch to primed microglia from other age-related changes, we correlated chromatin accessibility with the percentage of Micro2 cells in 16 donors, aged 46 to 82 (fig. S13), where age did not significantly correlate with Micro2 percentage (Pearson p-value = 0.082). We identified 858 cCREs with chromatin accessibility positively correlated with Micro2 percentages (“Primed state-correlated cCREs”, PCC > 0.6) (table S19), and 2,118 cCREs that positively correlated with donor age, despite cell state proportions (fig. S13, A to C, and table S20). While TF motif enrichment in the primed state-correlated cCREs were most significant for ETS and bZIP family TFs (Fig. 5M and table S21), age-correlated cCREs were most enriched for hormone-responsive elements of nuclear receptors (Fig. 5M), including glucocorticoid receptor (GR), which binds DNA at GR elements (GRE) in the presence of cortisol (*41*). Our results identify ETS and bZIP family TFs as putative drivers of epigenetic priming in microglia, while binding sites for hormone-regulated nuclear receptors such as GR have age-correlated activity independent of epigenetic priming.

### Correspondence Between 3D Genome Reorganization with the Epigenome and Transcriptome of Primed Microglia

Further leveraging our snm3C-seq data, we investigated 3D genome reorganization in primed microglia. The *IL15 gene* encodes a proinflammatory cytokine necessary for microglia activation (*42*) and displayed age-correlated expression (PCC = 0.79, FDR = 4.2e-6) (Fig. 6A). In Micro2, strengthened contacts were observed between the *IL15* promoter and an upstream cluster of cCREs with age-correlated chromatin accessibility and hypo-CG methylation (Fig. 6B), suggesting that these upstream cCREs are distal enhancers that drive expression of *IL15*. We also observed a region, centered over the promoter of *ZNF804A,* with several strengthened adjacent contacts in Micro2, forming a new local interaction domain (fig. S15B). *ZNF804A* showed the most significant age-correlated expression in microglia (fig. S15A), and harbors genetic risk variants for both schizophrenia and bipolar disorder (*43*). In addition, *ZNF804A* displayed age-correlated accessibility and hypo-CG methylation at its promoter and nearby cCREs (fig. S15B).

**Fig. 6.**
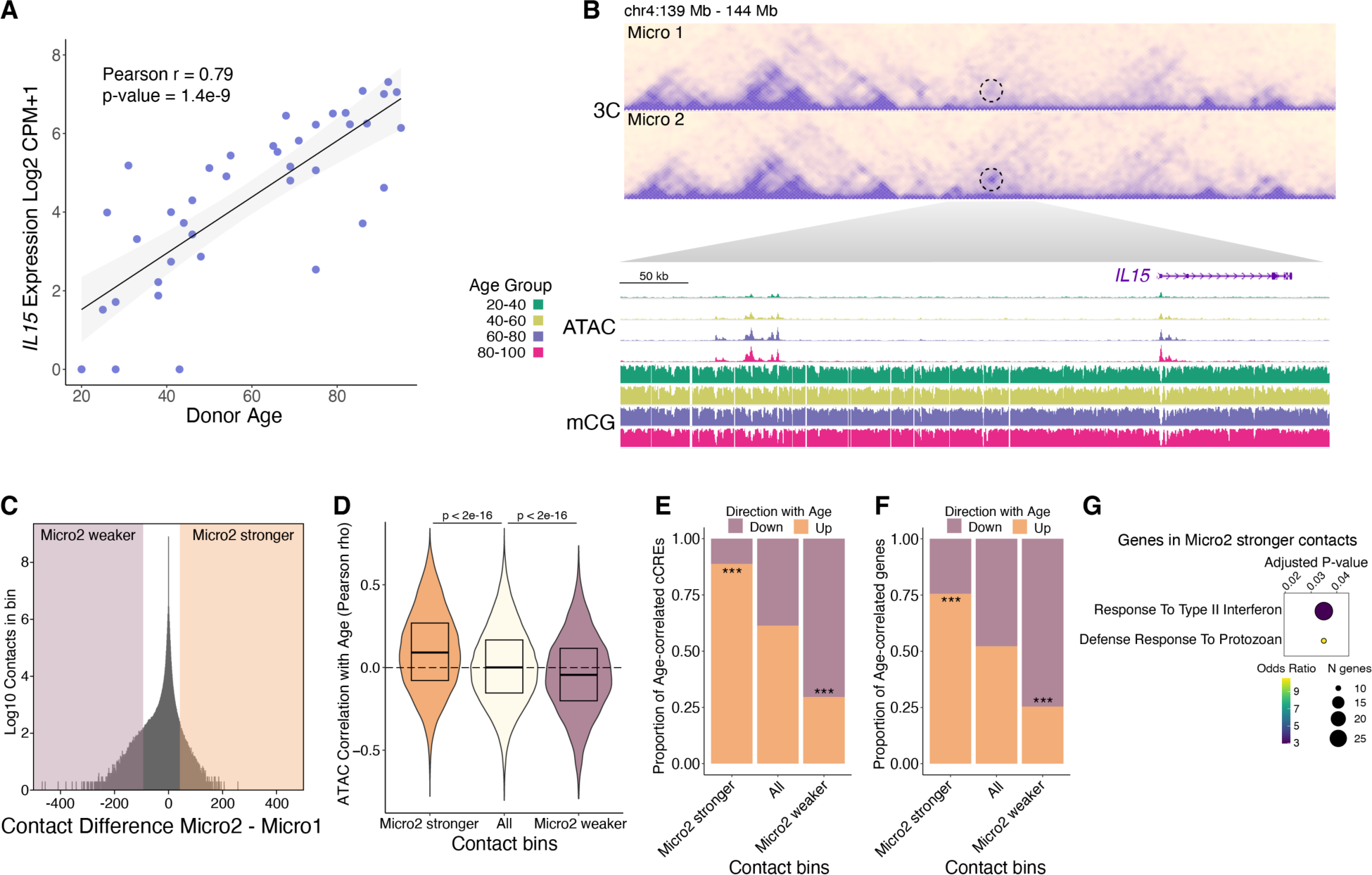
Correspondence between age-correlated 3D genome reorganization with the epigenome and transcriptome. (**A**) Scatter plot of *IL15* expression vs. donor age. (**B**) Heatmap showing contact frequency in Micro1 and Micro2. A dotted circle highlighting the contacting regions between the *IL15* promoter and an upstream cluster of cCREs. WashU browser snapshots of the circled region displaying chromatin accessibility and mCG fraction in microglia across age groups. (**C**) Density plot of contact differences, subtracting Micro2 from Micro1 contacts for each 100kb genomic bin after sampling equal number of total contacts for each group. Differential contact bins in the top ±0.0005% percentile are highlighted. (**D**) Violin plots showing the distribution of age-correlated chromatin accessibility for all microglia cCREs in the indicated contact bins. (**E** and **F**) Stacked bar plots show proportions of age-correlated cCREs (E) and age-correlated genes (F) within Micro2 differential contact bins. P-values computed using Pearosn’s Chi-square test comparing Micro2 higher up with All up or Micro2 lower down with All down, *** p < 0.001. (**G**) Gene ontology biological process enriched terms (q-value < 0.05) for genes having a transcription start site within a Micro2 higher contact bin.

Given these examples, we checked if 3D genome reorganization in primed microglia corresponded with epigenetic and transcriptomic changes, across the genome. We identified 100 kilobase (kb) genomic bins that had increased or decreased contact strength between Micro1 and Micro2 (table S22); referred to as “Micro2 stronger” and “Micro2 weaker”, respectively (Fig. 6C). Compared to all contact bins, the chromatin accessibility at cCREs in Micro2 stronger contact bins were more positively age-correlated, while chromatin accessibility at cCREs in Micro2 weaker contact bins were more negatively age-correlated (Fig. 6D). Because there can be many cCREs (500bp) within a 100kb contact bin, with most unlikely to be involved or influenced by any change in contact frequency, we focused only on age-correlated cCREs. We found an enrichment for age-correlated cCREs that go up with age in Micro2 stronger contact bins while Micro2 weaker contact bins were enriched with age-correlated cCREs that go down with age (Fig. 6E). We observed the same pattern of enrichment for age-correlated gene expression in Micro2 stronger and Micro2 weaker contact bins (Fig. 6F). The correspondence between age-related 3D genome changes with the epigenome and transcriptome was observed in additional glial types as contacts that are strengthened or weakened with age had similar trends of enrichment for age-correlated cCREs and genes (fig. S16, A and B).

Genes that had a transcription start site falling within a Micro2 stronger contact bin were functionally enriched for response to type II interferon (Fig. 6G), suggesting that 3D genome reorganization in primed microglia is preferentially occurring at the promoters of interferon-responsive genes. As with DNA methylation reprogramming, this conformational change in the 3D genome of Micro2 cells could reinforce a primed microglia state, where the chromatin is poised for a more powerful inflammatory response in the presence of immune challenges.

At the megabase scale, the genome is compartmentalized into two compartments, A and B, where chromatin contacts are largely constrained within each compartment (*44*). We observed a general weakening of compartmentalization in Micro2 compared to Micro1 cells and in the 80-100 age group (fig. S16, C to E). Since compartment A is enriched for euchromatin while compartment B is enriched for heterochromatin (*44*), we checked for correspondence between age-related changes in compartmentalization with the epigenome. We identified genomic regions with age-differential compartments in astrocytes, oligodendrocytes, and microglia, which were enriched for corresponding changes in accessibility with age (fig. S16F and table S23). Our results identify widespread age-related 3D genome reorganization coincident with age-correlated changes in the epigenome and transcriptome.

### Age-dependent Decay of Topologically Associating Domains and Decreased Accessibility at CTCF Binding Sites

We next extended our analysis to look at contacts between different chromosomes (trans contacts), which have been previously reported to change with age in mouse and human cerebellar granule cells (*9*). We observed a global increase in the number of trans contacts in the 80-100 age group for different cell types (fig. S14A), including microglia and oligodendrocytes (Fig. 7, A to D). Most cell types exhibited the increase in trans contacts but the change was cell type-dependent. For example, SUB excitatory neurons had increased trans contacts while CA and DG excitatory neurons had decreased trans contacts (fig. S14A), despite belonging to the same class.

**Fig. 7.**
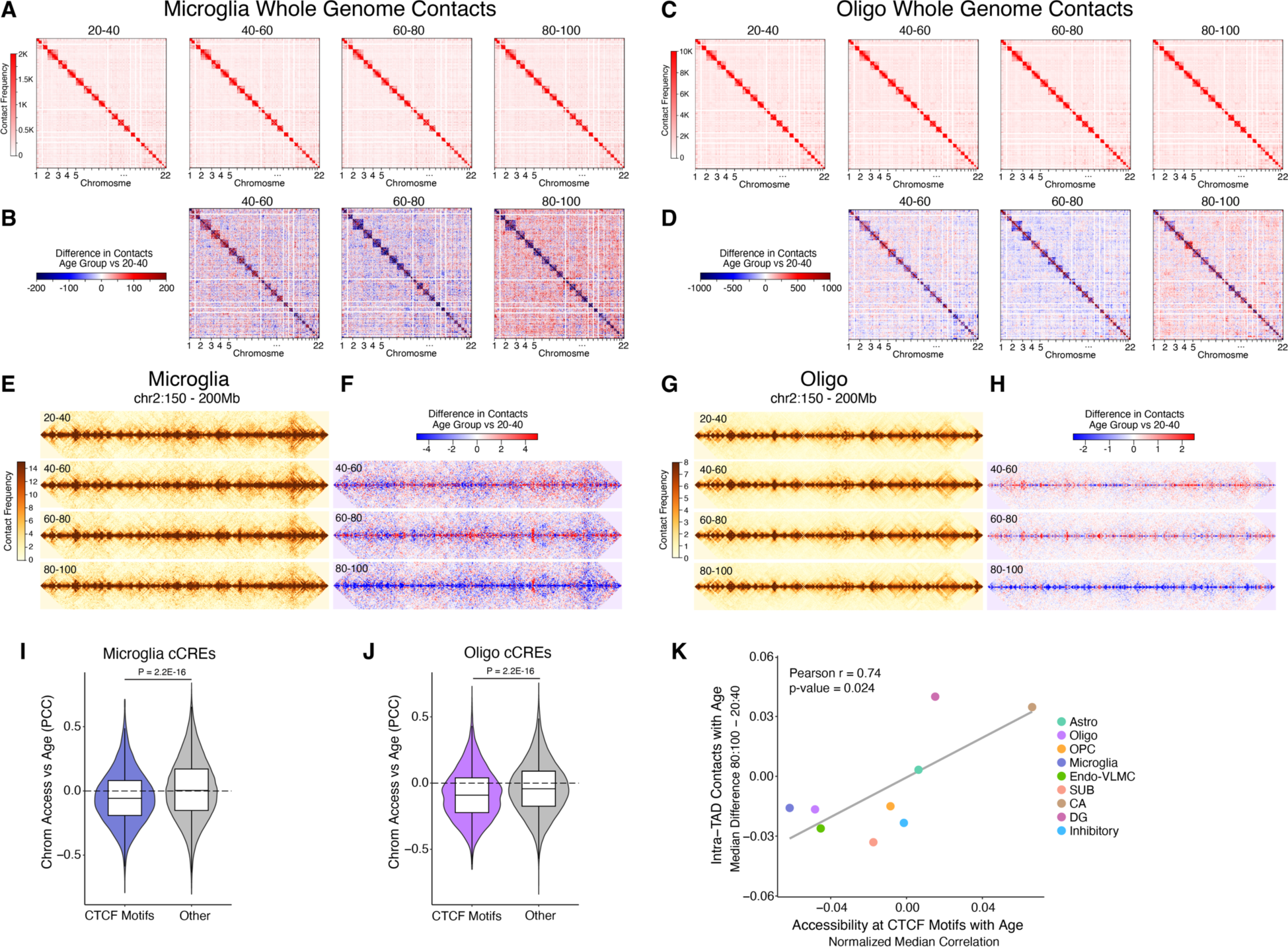
Global loss of homeostatic 3D genome architecture and CTCF binding in aging hippocampal cells. (**A**) Aggregated raw interchromosomal contact maps of microglia across different age groups. (**B)** Heatmaps showing the contact frequency differences in microglia between the 40-60, 60-80, and 80-100 age groups compared to the 20-40 age group. (**C**) Aggregated raw interchromosomal contact maps of oligodendrocytes across different age groups. (**D**) Heatmaps showing the contact frequency differences in oligodendrocytes between the 40-60, 60-80, and 80-100 age groups compared to the 20-40 age group. (**E**) Aggregated contact maps for microglia across each age group in the region chr2:150,000,000-200,000,000 at 100 kb resolution. (**F**) Heatmaps of contact frequency differences in microglia for the 40-60, 60-80, and 80-100 age groups compared to the 20-40 age group within chr2:150,000,000-200,000,000 at 100 kb resolution. (**G**) Aggregated contact maps for oligodendrocytes across each age group in the region chr2:150,000,000-200,000,000 at 25 kb resolution. (**H**) Heatmaps of contact frequency differences in oligodendrocytes for the 40-60, 60-80, and 80-100 age groups relative to the 20-40 age group within chr2:150,000,000-200,000,000 at 25 kb resolution. (**I**) Violin plots showing the distributions of age-correlation of chromatin accessibility at cCREs overlapping CTCF motifs or cCREs not overlapping CTCF motifs “Other” in microglia or (**J**) oligodendrocytes. (I and J) P-values derived from two-sided, unpaired Wilcoxon-rank sum test. (**K**) Scatter plot representing the difference in the median of all intra-TAD contacts (80:100 - 20:40 age groups) vs the median Pearson age-correlation of chromatin accessibility at CTCF motifs subtracted from the median Pearson age-correlation of all cCREs for each cell type.

Within chromosomes, chromatin is organized into self-interacting territories known as topologically associating domains (TADs) (*45*), where regulatory elements are largely constrained to regulate transcription within the same TAD. We found contacts within TADs were greatly diminished in the 80-100 age group (Fig. 7, E to H, fig. S14B, and table S24), specifically for cell types where trans contacts increased. This shift from intra-TAD contacts towards trans chromosomal contacts, highlights a widespread loss in homeostatic 3D genome organization in aging hippocampal cells.

CCCTC-binding factor (CTCF) is a master TF regulator of chromatin structure, and is critical for maintaining TADs (*46*). Therefore, we asked if the decay in TAD structures is a result of reduced CTCF DNA-binding in the aged hippocampal cells. In microglia and oligodendrocytes, chromatin accessibility at cCREs containing CTCF motif sequences was significantly more negatively correlated with age compared with cCREs not containing CTCF motifs (p-value < 2.2e-16) (Fig. 7, I and J). We next asked if CTCF DNA binding could explain cell type dependent decay of TADs by correlating loss of chromatin accessibility at CTCF motifs with TAD decay across cell types. Indeed, we found a strong correlation (Pearson r = 0.74, p-value = 0.024) between the change in intra-TAD contacts with chromatin accessibility at CTCF motifs with age (Fig. 7K), suggesting loss of CTCF DNA binding influences age-associated TAD decay. Although DNA methylation is known to block CTCF from binding (*47*), we did not observe an appreciable change in methylation at CTCF motifs with age (fig. S17), suggesting an alternative mechanism is responsible for the age-associated change in CTCF motif accessibility.

### Age-associated changes in neurons, oligodendrocytes, OPC, and endothelial cells

We also observed transcriptomic and epigenomic age-associated changes in cell types other than astrocytes and microglia, albeit fewer. We found that with aging, genes that function in axon guidance and synaptic potentiation had decreased expression in OPC and oligodendrocytes, and genes involved in maintenance of blood-brain barrier had decreased expression in endothelial cells (fig. S5B). Importantly, age-associated dysfunction in these pathways have been postulated to be key contributors to cognitive decline during aging (*15, 48*).

We also note the enrichment of binding sites for nuclear receptors, such as GR (NR3C1), at cCREs that gained chromatin accessibility with age in excitatory neurons (fig. S18A), oligodendrocytes, OPC, and microglia (fig. S6C). DNA binding activity of GR is stimulated by the presence of cortisol (*41*), which increases with age and elevated levels are associated with cognitive impairment and hippocampal atrophy (*49*). In age-decreased chromatin accessible cCREs of excitatory and inhibitory neurons, as well as oligodendrocytes (fig. S6C), we found enrichment for binding sites of CTCF (figs. S6C and S18, B to D), consistent with the observed age-associated changes to 3D genome organization (Fig. 7, and fig. S14).

## Discussion

In this study we conducted a comprehensive and in-depth investigation of epigenome and gene regulatory programs in the human brain during aging. We generated cell type-resolved transcriptomic, epigenomic, and 3D genomic profiles for different cell types from 40 hippocampus samples spanning the adult lifespan. Our age-balanced cohort empowered us to identify molecular features that change with age through robust age-correlation analyses. More specifically, we found genes whose expression increased or decreased with age in each cell type which revealed cell type-specific age-altered cellular processes, such as mitochondrial functions in astrocytes and neurons and inflammatory responses in microglia (Figs. 3C and 4B, and fig. S5, B and C). Additionally, we identified age-correlated changes in *cis*-regulatory element activities, allowing us to find putative pathways and transcription factor drivers of age-disrupted cellular processes in each cell type (Figs. 3F and 4D, and fig. S6C). Finally, we found age-differential 3D genome architecture which displayed high concordance with the aging epigenome and transcriptome (Fig. 6, and figs. S15 and S16, A, B, and F), highlighting the influence of gene regulatory features on gene expression during aging. Having a sex-balanced cohort, revealed that these gene regulatory features were affected in both females and males with age.

Surprisingly, we found a dramatic loss in the number of hippocampal astrocytes with age (Figs. 1, D and E, and 2, A and B, fig. S3, A, B, and E), which included a subset of astrocytes expressing tripartite synaptic markers (Fig. 2K). These astrocytes modulate pre- and postsynaptic neuronal functions through release of gliotransmitters after sensing neurotransmitters in the synaptic cleft (*17*). They are distinct in their expression of neurotransmitter sensing glutamate receptors, which in turn stimulate the release of gliotransmitters by increasing the intracellular calcium concentration (*17*). In addition to age-associated declines in astrocyte cell number, we found that astrocytes also had age-correlated loss of expression of genes critical for synthesizing ATP (Fig. 3C), a major gliotransmitter (*17, 18*). The functional implication of synaptic astrocyte loss is dysregulation of normal synaptic signaling which may directly contribute to cognitive decline such as impaired memory, as memories are thought to be initially stored in synaptic connections of the hippocampus (*50*). Beyond their synaptic roles, astrocytes are critical for blood brain barrier integrity through contacting the brain vasculature through their terminal processes (*51*). The dramatic loss of astrocytes in the aging hippocampus is likely to impact blood brain barrier permeability, which is known to increase with age in the human hippocampus (*52*).

While we could not determine the cause for astrocyte loss with age, our data point to potential mechanisms for their death that warrant further investigation. For example, loss of ATP synthesis can induce necrosis (*53*), a form of cell death distinct from apoptosis. Alternatively, ATP depletion can induce immune cell engulfment through exposure of phosphatidylserine on the outer leaflet of the plasma membrane— a so-called “eat-me” signal to phagocytosing cells (*54, 55*). Concurrent with down-regulation of ATP synthesis genes in aging astrocytes (Fig. 3C), we saw up-regulation of genes involved in lysosomal microautophagy (Fig. 3D), a cellular process intimately connected to cell death (*56*).

Through our single cell DNA methylome profiling, we discovered that in individuals between ages 50 and 75, hippocampal microglia undergo a cell state switch that involves reprogramming of their DNA methylomes (Fig. 5, A, B, D and F). Given the inter-individual heterogeneity of our cohort, along with the profiling of multiple molecular modalities, we were able to disconnect the cellular state composition from age, to identify putative drivers of this microglia state switch. We identified ETS family transcription factors as likely influencers (Fig. 5M and fig. S11D); however more investigation is necessary to uncover the mechanism inducing this age-related epigenetic switch to primed microglia.

The epigenetic reprogramming event results in DNA hypomethylation near genes that function in chemokine production, suggesting that these cells adopt an immune-activated microglia cell state (Fig. 5C). However, we did not see the same age-dynamic cell state switching when analyzing microglia transcriptomes. Therefore, this epigenetic state may not be associated with stable expression of proinflammatory genes, as many are cytotoxic and can only be transiently tolerated by cells (*57*). Instead, their DNA methylome reprogramming may reflect the primed immune state, where cells can quickly respond to a variety of insults, resulting in a rapid and elevated immune response. Indeed, mounting evidence suggests an age-dependent immune-sensitized state of hippocampal microglia (*26*). For example, aged rodents have heightened neuroinflammatory responses compared to young, specifically in the hippocampus, after various immune challenges including infections, surgery, or traumatic brain injury (*26*). These responses are associated with elevated levels of cytokines, including interleukin-1 beta, a proinflammatory gene that impairs memory when administered in young adult rats. While the source of this primed immune memory in aged hippocampal microglia has not been elucidated, an attractive notion is that it is faithfully encoded as DNA methylation, a relatively stable epigenetic mark (*58*).

Previously, it was shown that the 3D genome is reorganized over the lifespan in human and mouse granule cells (*9*) however, these changes did not coincide with changes in gene expression. Our study revealed a high degree of correspondence between 3D genome reorganization and transcriptomic and epigenomic dynamics with age (Fig. 6 and fig. S16, A, B, and F). In particular, primed microglia adopt a 3D genome conformation with strengthened chromatin contacts at genes related to type II interferon response (Fig. 6G), suggesting that, in addition to DNA methylome reprogramming, primed microglia adopt a 3D genome architecture amenable to heightened activation of inflammatory gene expression. Our data suggest a role for 3D genome reorganization in promoting neuroinflammation during human aging.

In most cell types, we observed a decay of TAD structures and a corresponding gain in trans-chromosomal contacts in the aged hippocampus (Fig. 7, A to H, and fig. S14). Loss of CTCF DNA binding, inferred through age-dependent decrease in chromatin accessibility at CTCF motifs, was observed in the same cell types exhibiting TAD decay with age (Fig. 7, I to K). As CTCF is an important regulator of hierarchical chromatin structure (*46*), its decreased DNA binding may be responsible for age-dependent loss of homeostatic 3D genome organization. Although we could not confidently attribute the global decay of TADs to changes in gene expression, it could have an influence on the fidelity of transcriptional programs, which may require more sensitive methods to measure.

## Acknowledgments

We acknowledge all members of the Ren and Xu labs for their helpful input.

## Author contributions

Conceptualization: BR, XX, CWC. Investigation: NRZ, NB, HSI, WMB, PKL, KD, BMG, AY, YT, CC, QZ, VA, LT. Formal analysis: NRZ, SL, SM, BY, BMG. Visualization: NRZ, SL, SM, BY, BMG, CS, DL, TW. Funding acquisition: BR, XX, CWC. Writing – original draft: NRZ, SL, SM, BR. Writing – review & editing: All authors contributed.

## Data and materials availability

Data produced in this study are available at NCBI GEO under accession number GSE278576.

## Supplementary Materials

Materials and Methods

Figs. S1 to S18

Tables S1 to S24

