## Supplemental Materials for "Epigenetic and 3D genome reprogramming during the aging of human hippocampus"

**The PDF file includes:**

Materials and Methods
Figs. S1 to S18

**Other Supplementary Materials for this manuscript include the following:**

Tables S1 to S24

### Materials and Methods

Nuclei preparation from frozen brain tissue for Chromium Single Cell Multiome ATAC + Gene Expression (10x Genomics)

Fresh frozen hippocampus tissue was obtained from 40 human donors postmortem. Tissues were pulverized in liquid nitrogen using a mortar and pestle on dry ice. Around 30mg of pulverized brain tissue was added to a ice-cold dounce homogenizer containing 1mL of chilled NIM-DP-L buffer (0.25M sucrose, 25mM KCl, 5mM MgCl2, 10mM Tris-HCl pH 7.5, 1mM DTT, 1X Protease Inhibitor (Pierce), 1U/μL Recombinant RNase inhibitor (Promega, PAN2515), and 0.1% Triton X-100). Tissue was dounce homogenized with a loose pestle (5-10 strokes) to shear large pieces, followed by a tight pestle (15-25 strokes) until the solution was uniform. Homogenate was filtered through a 30μm CellTrics filter (Sysmex, 04-0042-2316) into a LoBind tube (Eppendorf, 22431021) and pelleted (1000 rcf, 10 min at 4˚C) (Eppendorf, 5920 R). The pellet was resuspended in 1mL NIM-DP buffer (0.25M sucrose, 25mM KCl, 5mM MgCl2, 10mM Tris-HCl pH 7.5, 1mM DTT, 1X Protease Inhibitor, 1U/μL Recombinant RNase inhibitor) and pelleted (1000 rcf, 10 min at 4˚C). The pellet was resuspended in 400uL Sort Buffer (1mM EDTA, 1U/μL Recombinant RNase inhibitor, 1X Protease Inhibitor, 1% fatty acid-free BSA in PBS) with 2μM 7-AAD (Invitrogen, A1310). 120,000 7-AAD-positive nuclei were sorted (Sony, SH800S) into a LoBind tube containing Collection Buffer (5U/uL Recombinant RNase inhibitor, 1X Protease Inhibitor, 5% fatty acid-free BSA in PBS). Final collection volumes were determined and 5X Permeabilization Buffer (50mM Tris-HCl pH 7.4, 50mM NaCl, 15mM MgCl2, 0.05% Tween-20, 0.05% IGEPAL, 0.005% Digitonin, 5% fatty acid-free BSA in PBS, 5mM DTT, 1U/μL Recombinant RNase inhibitor, 5X Protease Inhibitor) was added to achieve a 1X solution. Nuclei were incubated on ice for 1 minute, then centrifuged (500 rcf, 5min at 4C) in a swinging bucket centrifuge. Supernatant was discarded and 650uL of Wash Buffer (10mM Tris-HCl pH 7.4, 10mM NaCl, 3mM MgCl2, 0.1%.Tween-20, 1% fatty acid-free BSA in PBS, 1mM DTT, 1U/μL Recombinant RNase inhibitor, 1X Protease Inhibitor) was added without disturbing the pellet followed by centrifuging (500 rcf, 5 min at 4˚C) in a swinging bucket centrifuge. Supernatant was removed, and the pellet was resuspended in 7uL of 1X Nuclei Buffer (Nuclei Buffer (10x Genomics), 1mM DTT, 1 U/μL Recombinant RNase inhibitor). 1μL was used for counting on a hemocytometer after staining with Trypan Blue (Invitrogen, T10282). 16-20k nuclei were used for tagmentation reaction and 10x Genomics controller loading. Then libraries were generated following the manufacturer's recommended protocol (https://www.10xgenomics.com/support/single-cell-multiome-atac-plus-gene-expression). 10x multiome ATAC-seq and RNA-seq libraries were paired-end sequenced on a NextSeq 2000 to assess data quality. If data quality was satisfactory, libraries were deep sequenced on a NovaSeq 6000 to a target depth of ~50,000 reads per cell for each modality.

10x multiome sequence data processing and clustering

Raw sequencing was processed using cellranger-arc v2.0.0 (10x Genomics), generating snRNA-seq UMI count matrices for intronic and exonic reads mapping in the sense direction of a gene for hg38 GRCh38 human genome assembly and Gencode v32 gene annotations. We performed unsupervised clustering with RNA UMI counts using Seurat v5 (*59*) integrative analysis pipeline. First, cell barcodes from the raw cellranger matrices were filtered for low quality nuclei by requiring ≥ 500 ATAC fragments, ≥ 200 genes detected per nuclei, and < 20% mitochondrial RNA reads. We further required > 5 TSS enrichment (GRCh38 GENCODE v46) using SnapATAC2 (*60*). Counts were normalized using SCTransform and 2,000 protein coding variable genes were determined and used for principal component analysis (PCA). Putative multiplets were predicted using DoubletFinder (*61*) software and 10% of cell barcodes were removed from each sample that had the highest doublet score. Batch correction across donors was performed using reciprocal principal component analysis (rPCA) on SCTransformed PCs. A k-nearest neighbors graph was built using PCs 1:30 and clusters were identified using leiden clustering. To visualize clusters, we performed the non-linear dimension reduction technique, UMAP (*62*). We annotated subclass-level cells by known marker gene expression and reference mapping to a published hippocampus snRNA-seq dataset (*63*, *64*) using Seurat. Additional multiplet clusters were identified and removed for expressing multiple cell type-specific marker genes.

<u>ATAC-seq peak calling and filtering</u>

For each cell type, we divided the cells into four donor age groups, 20-40, 40-60, 60-80, and 80-100 years. For each cell type, we called peaks using ATAC-seq fragments of combining cells from all 40 donors, as well as in each of the four individual age groups, to capture any age-specific peaks. This gave us 5 lists of peaks for each cell type. We called peaks using MACS2 (*65*) software with the following command: macs2 callpeak --shift -75 --ext 150 --bdg -q 0.1 -B --SPMR --call-summits -f BAMPE. We extended each peak summit by 249 bp upstream and 250 bp downstream to achieve a uniform 500 bp span for each peak. Peaks overlapping with ENCODE blacklist regions for the hg38 genome assembly (https://mitra.stanford.edu/kundaje/akundaje/release/blacklists/) were removed. To account for variability in sequencing depths because of differences in cell type abundance, MACS2 peak scores (-log10(q-value)) were converted to 'score-per-million' (SPM). We retained peaks with a score-per-million ≥ 4 in any of the 5 peak sets for each cell type. We then created a union set of peaks across all cell types and age groups by merging all peaks that have any overlap, then resized to 500bp using the peak summit with the highest SPM from any individual peak set. From this union set, we filtered out peaks that did not meet the score-per-million ≥ 4 threshold in any cell type, ensuring that all retained peaks were robustly detected in at least one cell type. To reduce redundancy, coordinates for peaks of each cell type were replaced with coordinates from the refined union set of peaks, by intersecting the filtered peaks from each cell type with the refined union set using bedtools intersect (https://bedtools.readthedocs.io/en/latest/content/tools/intersect.html).

<u>Nuclei isolation and Fluorescence Activated Nuclei Sorting (FANS)</u>

Fresh frozen hippocampus tissues were pulverized in liquid nitrogen using a mortar and pestle on dry ice. Around 20mg of pulverized brain tissue was added to a ice-cold dounce homogenizer containing 1mL of chilled NIM-DP-L buffer (0.25M sucrose, 25mM KCl, 5mM MgCl2, 10mM Tris-HCl pH 7.5, 1mM DTT, 1X Protease Inhibitor (Pierce), 1U/µL Recombinant RNase inhibitor (Promega, PAN2515), and 0.1% Triton X-100). Tissue was dounce homogenized with a loose pestle (5-10 strokes) to shear large pieces, followed by a tight pestle (15-25 strokes) until the solution was uniform. Homogenate was filtered through a 30µm CellTrics filter (Sysmex, 04-0042-2316) into a LoBind tube (Eppendorf, 22431021) and pelleted (1000 rcf, 10 min at 4˚C) (Eppendorf, 5920 R). Pellets were resuspended in 2% formaldehyde in DPBS for 5 minutes at room temperature on a rotator. Glycine was added to obtain 0.2M glycine and tubes were incubated for 5 min on a rotator at room temperature. Lysate was centrifuged at 1,000 x g for 10 min at 4°C (in bucket rotor). Supernatant was removed and the pellet was resuspended in 1mL of cold DPBS. Lysate was centrifuged at 2,500 x g for 5 min at 4°C (in bucket rotor). Supernatant was removed and the pellets were used for in situ 3C following the steps in the Arima-3C Single Cell Beta Kit (Arima Genomics), with some modifications. Briefly, 44uL of Conditioning Solution was added, and lysate was transferred to a 0.2 mL PCR tube, and mixed gently by pipetting. Tubes were incubated at 62°C for 30 min in a thermal cycler, with the lid temperature set to 85°C. 20µL of Stop Solution 2 was added, and mixed gently by pipetting. Tubes were incubated at 37°C for 15 min in a thermal cycler with a lid temperature set to 85°C. 28µL of the restriction enzyme digestion mix (9.8uL water, 9.2uL Buffer H, 4.5uL Enzyme H1, 4.5uL Enzyme H2) was added and gently pipette mixed. Tubes were incubated in a thermal cycler for 37°C for 60 min, 65°C for 20 min, then 25°C for 10 min. 82uL of ligation mix was added (70uL Buffer C, 12uL Enzyme C) and gently mixed by pipetting, then incubated at 25°C for 15 min. Lysate was transferred to a tube containing 850µL 1% BSA in DPBS on ice and filtered through a 30um Celltrics strainer to a new tube on ice. Lysate was centrifuged at 1,000 x g for 10 min at 4°C (in bucket rotor). Supernatant was removed and pellets were resuspended each sample in 800uL of 1% BSA in DPBS. Nuclei were stained with DRAQ7 at 1:200 and mixed gently with a pipette and incubated on ice for 5 min. One tube of Proteinase K (Zymo cat. no. D3001-2-D) was dissolved in 1mL of Proteinase K Resuspension Buffer (Zymo cat. no. D3001-2-B) then added to a mix of 15mL M-Digestion Buffer 2x (Zymo cat. no. D5021-9) and 14mL water to make pK Digestion Buffer. 2uL of pK Digestion Buffer were added to each well of an Eppendorf twin.tec® PCR Plate 384 LoBind®, skirted, 45 µL, PCR clean (Eppendorf cat. No. 0030129547). Single DRAQ7 positive nuclei were sorted into individual wells of the 384-well plate using a Sony SH800 cell sorter. Plates were sealed then incubated in a thermal cycler at 50°C for 20 min with a heated lid at 85°C for nuclei digestion.

Library preparation and Illumina sequencing for snm3C-seq

The snm3C-seq samples followed the library preparation protocol detailed previously (*10*, *66*). This protocol has been automated using the Beckman Coulter Biomek i7 liquid handlers for reactions in 384 and 96-well plates. The snm3C-seq libraries were shallow sequenced on the Illumina NovaSeq 2000 to 2-10M reads to check quality. Successful libraries were deeply sequenced on the NovaSeq X Plus instrument, using one lane of 25B flow cell per 4 384-well plates and using 150 bp paired-end mode.

Preprocessing of snm3C-seq

The snm3C-seq FASTQ reads were demultiplexed and mapped using YAP (Yet Another Pipeline) software (cemba-data v1.6.9), as previously described (*66*). First, FASTQ files were demultiplexed for each cell barcode. Next, two-pass mapping was performed with bismark (v0.20), using bowtie2 (v2.3). BAM file processing and QC was performed using samtools (v1.9) and picard (v3.0.0). Chromatin contacts were called and methylome profiles were generated using Allcools (v1.0.23). All reads were mapped to the hg38 reference genome assembly.

Quality control of snm3C-seq

Low quality cells were removed by requiring Bismark mapping rate for both reads R1 and R2 > 0.5; total final reads > 100,000; mCCC level < 0.03; mCH level < 0.2; mCG level > 0.5. Potential multiplets were identified using the MethylScrublet function in AllCools software for each sample, and cells with a doublet score > 0.015 were removed. Further quality filtering involved removing a low-quality cluster. Specifically, a cluster consisting of 36 cells with hypo-methylation of mixed cell type markers was excluded. Finally, 22,240 cells were included in the downstream analysis.

Methylome Clustering Analysis

For the cells passing quality control, we generated the methylome cell dataset (MCDS) from single-cell DNA methylome profiles of the snm3C-seq datasets using "allcools generate-dataset" command. This dataset comprises a cell-by-feature tensor that encompasses various genomic regions and types of DNA methylation contexts. We used hypo-methylation probability (hypo-score) in CGN contexts across non-overlapping 5kb bins (chrom5kb) on the hg38 chromosome. And we used gene body regions ±2 kb for clustering annotation and integration with the 10x multiome dataset. We performed clustering by (1) binarizing the matrix; (2) excluding 5kb bins that had a non-zero methylation level in below 10% of the cells; (3) normalizing using term frequency–inverse document frequency (TF-IDF); and (4) reducing dimensionality using truncated singular value decomposition (SVD). Then, we applied batch balanced k-nearest neighbor (BBKNN) (*67*) to remove technical batch effects, followed by visualization using UMAP (*62*). Finally, we performed graph clustering using the Leiden algorithm (*68*).

Cluster-level DNA Methylome Analysis

After methylome clustering analysis, we merged the single-cell ALLC files into pseudo bulk level using the "allcools merge-allc" command. Next, we performed DMR calling using 'call_dms' and 'call_dmr' functions in AllCools. In brief, we calculated CpG differential methylated sites (DMS) using a permutation-based root mean square test (*69*). The base calls of each pair of CpG sites were added before analysis. We then merged the DMS into DMR if they are (1) within 500 bp and (2) the minimum methylation difference was greater than or equal to 0.3 across samples. We applied the DMR calling framework across the cell clusters and cell clusters for each age group.

Chromatin contact matrix and imputation

After chromatin contact data were generated by the YAP pipeline, we used the cis-long range contacts (contact anchors distance > 1,000 bp) and trans contacts to generate single-cell chromatin contact matrices without imputation (hereafter referred to as raw contact matrices) at three genome resolutions: chromosome 100-Kb resolution for the chromatin compartment analysis; 25-Kb bin resolution is for the chromatin domain boundary analysis; and 10-Kb resolution. The raw cell-level contact matrices for all resolutions are stored in HDF5-based mcool format. We then used the scHiCluster package (v1.3.5) to perform contact matrix imputation. In brief, the scHiCluster imputes the sparse single-cell matrix in two steps: the first step is Gaussian convolution (pad=1); the second step is to apply a random walk with restart algorithm on the convoluted matrix. The imputation is performed on each chromosome? of each cell. For 100-Kb matrices, the whole chromosome is imputed; for 25-Kb matrices, we imputed contacts within 10.05Mb; for 10-Kb matrices, we imputed contacts with 5.05Mb. The imputed matrices for each cell were stored in cool format. For most following analyses, cell matrices were aggregated into cell groups identified in the previous section and stored in cool format.

10x multiome and snm3C-seq single cell integration

The mCG fraction of gene body regions ±2 kb ("hypomethylation gene score") was used for integration with the 10x multiome snRNA-seq dataset. The snRNA-seq data were normalized by dividing the counts of each cell by the total counts across all genes. And logarithmic transformation was applied to both datasets. Highly variable genes and gene body regions ±2 kb were identified independently from both snRNA-seq and snm3C-seq datasets, and the datasets were scaled. Before integration, the mCG fraction was multiplied by -1 to account for the negative correlation between DNA methylation level and gene expression. Following integration, dimensional reduction was performed based on the shared set of highly variable genes. Harmony (*70*) was applied for a batch corrected embedding. Finally, cell type labels from the 10x multiome dataset were transferred to predict cell types in the snm3C-seq dataset using a Euclidean nearest neighbor descent approach implemented in PyNNDscent (*71*).

Age-correlation of genes, cCREs, and DMRs

To identify genes, cCREs, or DMRs with age-correlated activity in each cell type, we calculated Pearson correlation coefficients for pseudo bulk CPM (gene expression or chromatin accessibility) or mCG fractions (DMRs) vs donor age. Due to differences in cell numbers captured for each donor from a given cell type, we filtered donors and features, using different criteria for each modality. For gene expression, we removed donors that had less than 20,000 total RNA counts. For the remaining donors, we removed genes with low expression by requiring the total counts across remaining donors must be equal or greater than 2 multiplied by the number of remaining donors. For example, in cell type *n* if there are 30 donors remaining after filtering, genes must have a total count of 60 or greater when counts are summed across these donors. For chromatin accessible cCREs, all cCREs identified in a given cell type were considered. Donors with less than an average of 1 count per number of cCREs, were removed from the correlation analysis. All DMRs identified in a given cell type were considered. Donors with less than an average of 1 count per number of DMRs, were removed from the correlation analysis. Pearson correlation p-values were converted to FDR adjusted p-values. Features with a FDR < 0.1 were considered for downstream analysis.

Astrocyte sub-clustering

We employed an iterative clustering approach to identify astrocyte subclusters. We subsetted the astrocytes and re-normalized using SCTransform, identifying 3,000 highly variable genes. After scaling, we performed principal component analysis (PCA) and used Harmony for batch correction. We then utilized the first 30 dimensions of the Harmony-corrected data to find neighbors and generate a UMAP visualization. To determine the optimal resolution for clustering, we implemented an iterative clustering method as described in Li et al[13]. We then used resolution 0.2 for sub clustering and found 7 subclusters. To identify putative tripartite astrocyte clusters, we calculated module scores using Seurat's AddModuleScore function for 18 tripartite synaptic genes (*19*) *NRXN1*, *NLGN3*, *ITGB1*, *ITGAV*, *CDH2*, *CADM3*, *CADM1*, *NCAM1*, *TGFB2*, *ATP1B2*, *EPHA4*, *GJB6*, *BCAN*, *CSPG5*, *SPARCL1*, *SDC2*, *SDC4*, *GPC1*.

Differential gene expression and chromatin accessibility analysis

We used MAST (*72*) on single cell normalized counts to identify differential expressed genes and chromatin accessible cCREs between cell clusters, namely Astro1 vs Astro2 and Micro1 vs Micro2. For Astro1 vs Astro2 differentially expressed genes, we used donor as a latent variable. We required 10% of cells to be expressing the gene, p-adjusted < 0.05, and log2 fold change > 1. For Astro1 vs Astro2 differential cCREs, we called peaks on astrocytes and then employed logistic regression as recommended in Signac for differential peak analysis. We used donor as a latent variable. We required 10% of cells to have the cCRE accessible, p-adjusted < 0.01. For Micro2 and Micro1 differential gene expression we used age as a latent variable. We considered genes expressed in 1% of cells and p-adj < 0.0001. For Micro1 and Micro2 differential cCREs, we considered cCREs accessible with p-adj < 10e-24 due to inflated p-values and to obtain an appropriate number of elements for downstream analysis.

Gene ontology enrichment analysis

We performed GO enrichment analysis using the GSEApy (*73*) and ClusterProfiler (*74*). For each gene set, we used GO biological process terms. We performed such analyses using the most appropriate background set: for astro and microglia subcluster marker genes, we used genes expressed in 10% of cells, and for age-correlated genes, we used genes tested for correlation after filtering low count genes (see above). For GO enrichment analysis for DMRs, we first identified DMRs using the 'call_dmr' function in AllCools and filtered the DMRs to include those with more than 3 DMSs. Next, we used GREAT (*75*) with default settings to assign DMRs to genes that were hypo-methylated in either Micro1 or Micro2. GO enrichment analysis for biological process was performed using GREAT with a background gene set consisting of all DMRs in microglia.

Transcription factor motif enrichment analysis

For transcription factor (TF) motif enrichment analysis, we utilized the HOMER database. We focused on peaks that correlated with age, using significant peaks with a false discovery rate (FDR) of 0.1. As background, we employed peaks accessible in the specific cell type of interest. The enrichment analysis was performed using these parameters, and we considered only statistically significant TFs in our results. This approach allowed us to identify age-related changes in TF binding patterns within our dataset. To assess transcription factor (TF) activity at the single-cell level in excitatory or inhibitory neurons, we employed ChromVAR (*76*). For identifying differential TF activity, we utilized the FindMarkers function from the Seurat package, applying it to the ChromVAR z-scores. We set the mean.fxn parameter to rowMeans and fc.name to "avg_diff" for this analysis. To focus on the most relevant and robust results, we applied the following filtering criteria: 1) TFs showing activity in at least 25% of cells, 2) adjusted p-value < 0.05, 3) minimum difference of 10% in activity between old and young samples.

Immunofluorescent staining

Immunolabeling was conducted on 30µm-thick free-floating hippocampal coronal sections. Tissues underwent sodium citrate (BioLegend, San Diego, CA; #928502) antigen retrieval for 30 minutes at 90°C and were subsequently washed three times with 1x phosphate buffered saline (PBS) for 5 minutes each after returning to room temperature. Sections were then bathed in lipofuscin autofluorescence blocking TrueBlack solution (Biotium, Fremont, CA; #23007) for 5 minutes, rinsed three times with 1xPBS, and incubated in blocking buffer containing 5% normal donkey serum (Jackson ImmunoResearch Laboratories, West Grove, PA; #017-000-121) and 0.075% (v/v) Triton X-100 in 1xPBS for 2 hours at room temperature. Brain slices were then incubated in blocking solution containing an antibody against astrocytic marker ALDH1L1 (LSBio, Shirley, MA; #LS-C796716; 1:200) for 24 hours at 4°C. Sections were then rinsed three times with 1xPBS and incubated for 2 hours in blocking solution containing Alexa Fluor 488-conjugated donkey anti-mouse secondary antibody (Jackson ImmunoResearch Laboratories, West Grove, PA; #715-545-150) and DRAQ5 nuclear stain (ThermoFisher, Waltham, MA; #62254; 1:2,000). The tissues were then rinsed with 1xPBS and finally mounted and coverslipped with Fluoromount-G (SouthernBiotech, Birmingham, AL; #0100-01) for microscopy.

Brain section imaging and quantification

Hippocampal CA3 confocal micrographs were obtained using an Olympus FV3000 microscope. Serial optical sections were captured for each brain slice using a 20x objective with a 2x zoom and a 0.89 μm step interval for a total of 10 slices (z-stack). All images were acquired using identical conditions. Maximum intensity projections from z-stacks were processed using the Olympus FV31S-SW Fluoview software (Version 2.6, Olympus Life Science). The resulting 2D projection images (0.101 $mm^2$ ROI area for each) were analyzed using Photoshop. ALDH1L1-positive astrocytes were manually counted in hippocampal CA3 from young (n=5; 20–38 y.o.) and aged (n=7; 80–106 y.o.) donor groups. Values were normalized by the number of DRAQ5 nuclei and statistically evaluated using an unpaired Welch's T-test & visualized using GraphPad Prism 9 (GraphPad Software, CA, USA). For microglia soma size quantification, three CA3 confocal micrographs were taken from each of the young (n=3; 20-38 y.o.) and aged (n=6; 80-106 y.o.) donor's hippocampus. IBA1+ microglia 2D projection images (0.0262 mm2 ROI area) were then analyzed using ImageJ/FIJI software. The polygon selection tool was used to manually outline microglia somas and later quantified using the measure tool. The resulting data was statistically analyzed using a linear mixed-effect model (LME) to address the issue of correlated data arising from repeated measurements from single donors (*77*). LME analysis was performed on the average microglia soma size from each ROI (3 per donor) using the MATLAB "fitlme" function.

Identifying putative enhancer–gene pairs with the ABC model

We used the Activity-by-Contact (ABC) model (*29*) to identify putative enhancers and target genes in each cell type. In brief, the ABC model uses normalized contact frequencies from Hi-C data, along with a measure of enhancer activity, to predict putative enhancer–gene pairs. For each cell type, we provided normalized Hi-C matrices as bedpe files at 10 kb resolution, ATAC-seq BAM files for each cell type and a list of cCREs from ATAC-seq peaks identified in that same cell type. Only genes expressed with CPM 1 or greater were considered. Predictions were considered positive and used for downstream analyses if ABC score was 0.02 or greater.

Compartment analysis

We used "hicluster compartment" command in scHicluster v1.3.5 (*78*) to compute compartment score using the cluster-level imputed contact matrices at 100 kb resolution contact matrices obtained after down-sampling cells. The down-sampling was performed to ensure a comparable total number of contacts for pseudo bulk analysis, which was carried out per age group within each cell type.

Differential contacts and differential compartments

We used raw contact matrices to look for chromatin contacts in *cis* with large differences between age groups or subtypes. To remove the baseline bias introduced by differences in sequencing depth between conditions under comparison, we downsampled the total number of contacts in each condition to the same level and regenerated the raw contact matrices. For each comparison, we extracted the *cis* contacts within the top ±0.0005% percentile from the distribution of differences in coverage between the conditions. To identify the differential compartments, we used dcHiC (*79*). First, raw PC values were generated for each chromosome and group, followed by quantile normalization of these PC values. Subsequently, for each 100 kb bin, Mahalanobis distance, a multivariate z-score measuring the degree to which each bin deviates as an outlier from the overall compartment score distribution across all datasets, was computed and used to determine statistical significance through FDR correction. A compartment was considered differential if its adjusted p-value was below 0.05.

Intra-TAD contact quantifications

We used scHicluster v1.3.5 (*78*) (hicluster domain) to identify topologically associated domains (TADs) on downsampled raw contact matrices at 10 kb resolution. We counted the number of contacts by summing up the contacts within the boundaries of each TAD for every single cell. These intra-TAD contacts were divided by the number of cis long contacts + trans contacts for each cell to obtain the ratio of intra-TAD contacts in each age group or microglia subcluster.

DNA Methylation Aging Clock

To predict biological age using DNA methylation, we calculated mCG fractions at the 353 CpG sites listed in the Horvath epigenetic clock (*31*) using "allcools allc-to-region-count" command. We employed the pyaging tool (*83*) for age prediction, utilizing KNN as an imputation strategy for missing CpG ratios. Biological age predictions were performed both for each cell type separately and for all cell types combined (whole tissue). We calculated Pearson correlation coefficients between chronological and predicted biological age to address data sparsity in cell types with lower abundance, we computed pseudo donor methylation ratios after grouping data from 2, 4, 5, or 8 donors with adjacent chronological ages, subsequently calculating their predicted biological age. Average chronological age for binned donors was used for correlation with predicted biological age. This approach allowed us to mitigate the impact of missing data points to still evaluate biological age predictions for cell types with lower abundance.

External dataset

NRF1 ChIP-seq in neuroblastoma cell line SK-N-SH signal p-value bigwig (Accession ENCFF762LEE) was downloaded from the ENCODE portal (https://www.encodeproject.org/experiments/ENCSR000EHZ/).

Data Availability

Data produced in this study are available at the NCBI GEO under accession number GSE278576.

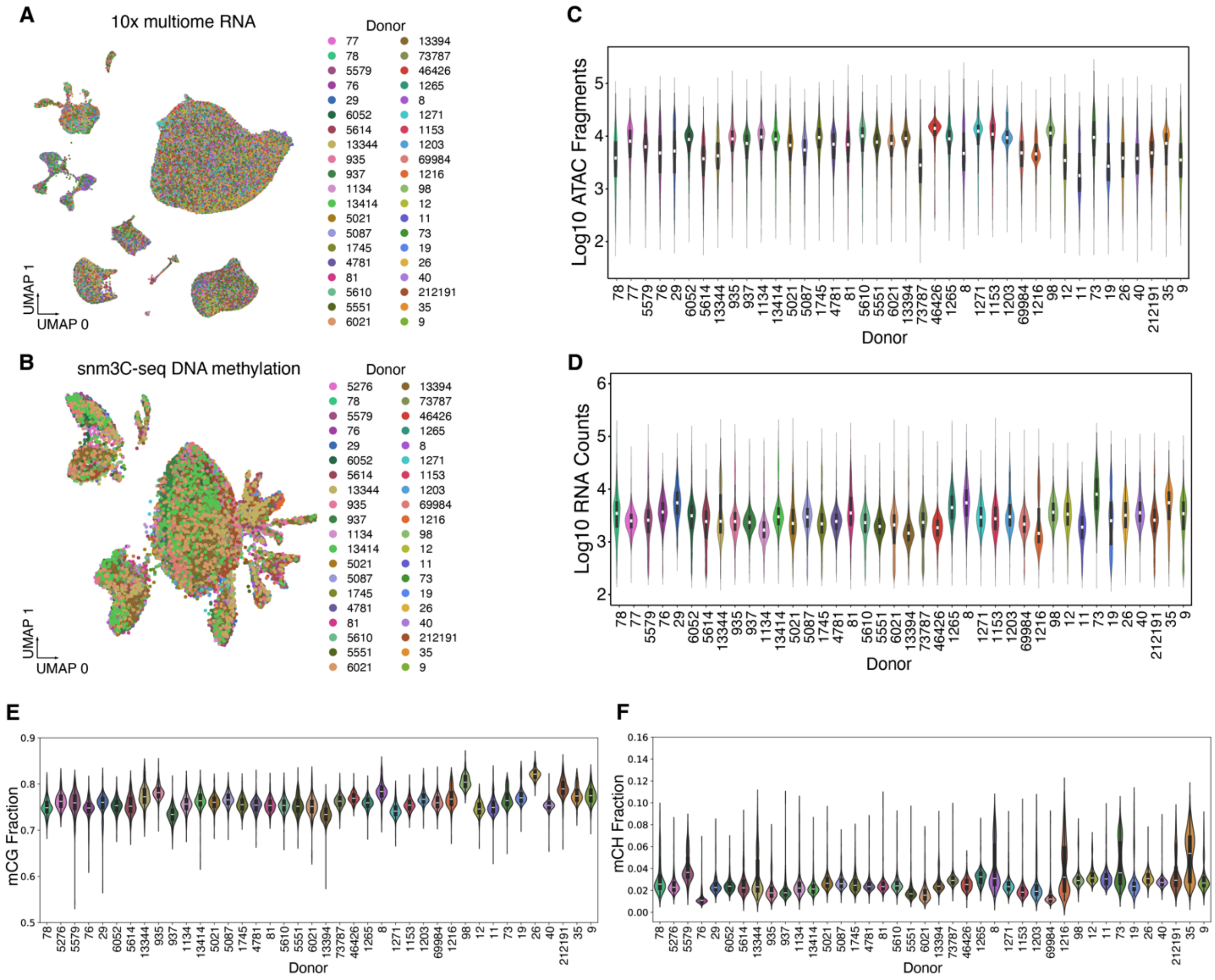


**Fig. S1.**
10x multiome and snm3C-seq quality metrics across samples. (**A** and **B**) UMAP embeddings of 10x multiome RNA (A) and snm3C-seq DNA methylation (B) clustering, colored by donor (n=40). (**C** and **D**) Violin plots showing distribution of log10 ATAC-seq fragment counts (C) and log10 RNA read counts (D) for each donor. (**E** and **F**) Violin plots showing distribution of snm3C-seq DNA methylation mCG fraction (E) mCH fraction (F) for each donor.

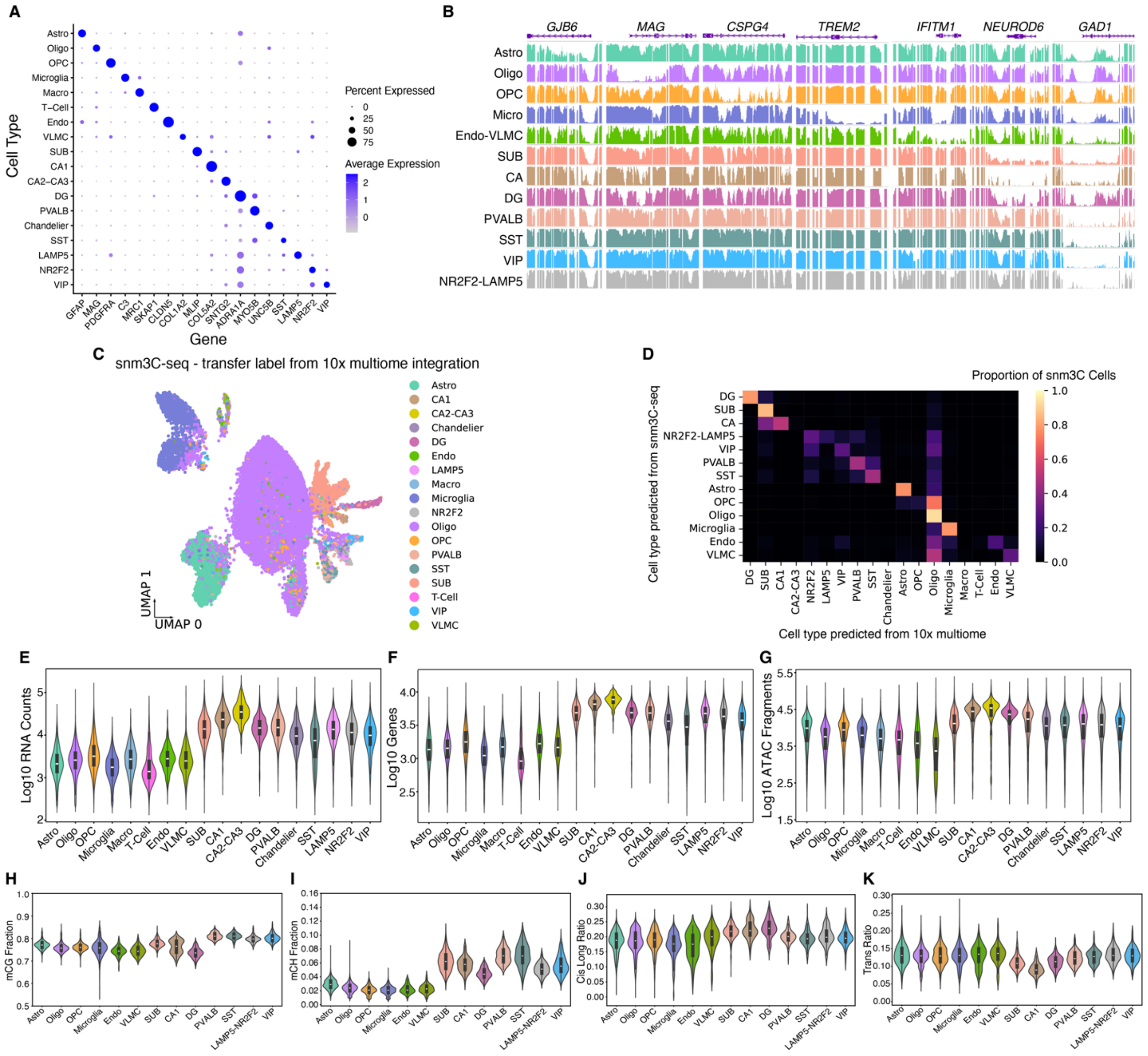

**Fig. S2.**

Single cell annotations and feature metrics. (**A**) Dot plot showing the average expression and proportion of cells expressing cell type marker genes across cell types. The color and size of each dot indicate the expression level and percentage of cells expressing genes, respectively. (**B**) mCG fraction in cell type marker gene regions at the pseudo bulk level for each cell type. (**C**) UMAP embedding of snm3C-seq DNA methylation data, annotated with cell type labels transferred from 10X multiome RNA data using hypomethylation gene scores and RNA integration. (**D**) Heatmap showing the proportion of snm3C-seq cells compared between annotations based on snm3C-seq data alone and labels transferred from 10X multiome RNA data. (**E** to **G**) Violin plots showing distribution of log10 RNA-seq read counts (E), log10 number of genes (F), and log10 ATAC fragments (G) for each cell type. (**H** to **K**) Violin plots showing distribution of mCG fraction (H), mCH fraction (I), cis-long ratio (J), and trans ratio (K) for each cell type.

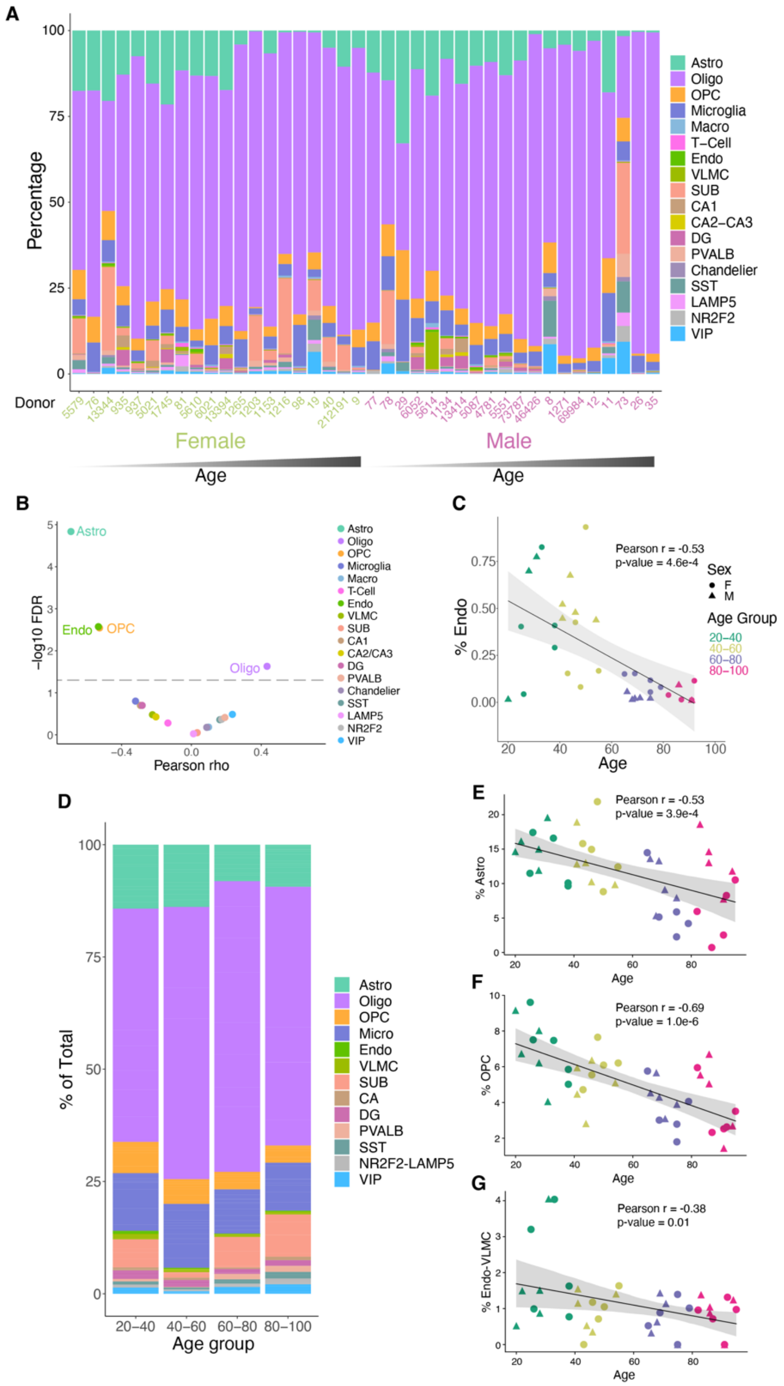


**Fig. S3.**
Cell type proportional changes with age. (**A**) Stacked bar plots representing percentage of cell type for each donor from 10X multiome RNA-seq. (**B**) Scatter plot showing the Pearson correlation coefficients (rho) of age vs negative log10 FDR. Each dot represents a cell type. (**C**) Scatter plots showing the percentage of endothelial cells vs donor age recovered from 10x multiome experiments. (**D**) Stacked bar plots representing percentage of cell type recovered from donors in indicated age group from snm3C-seq data. (**E** to **G**) Scatter plots showing the percentage of astrocytes (E), OPCs (F), and endothelial cells and VLMCs (G) vs donor age recovered from snm3C-seq experiments.

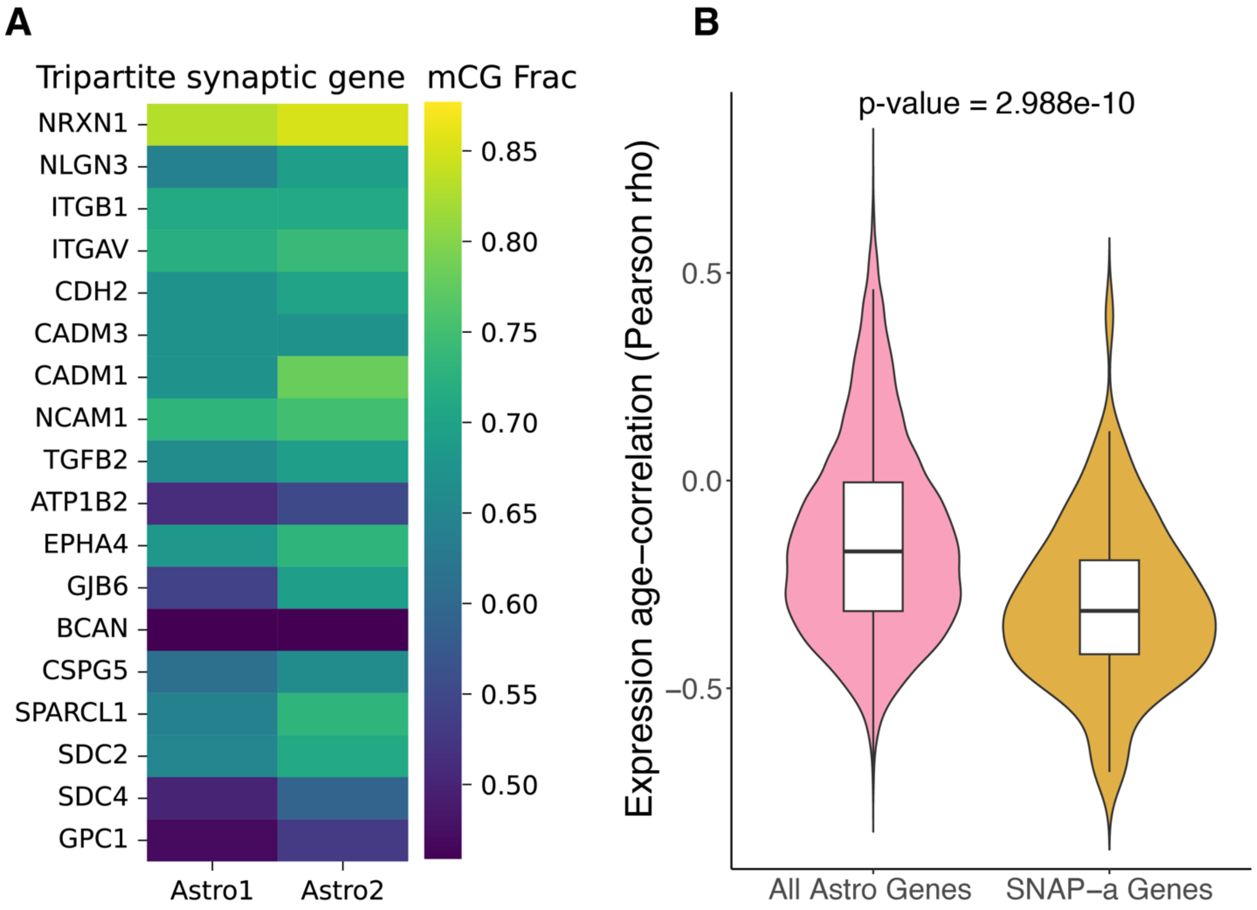


**Fig. S4.**

Synaptic astrocyte gene DNA methylation and gene expression. (**A**) Heatmap showing the mCG fraction of 18 tripartite synaptic genes (*19*) in astrocyte subclusters from snm3C-seq data. (**B**) Violin plots of astrocyte gene expression of all expressed genes and synaptic astrocyte genes reported to decrease in aging human prefrontal cortex (*23*).

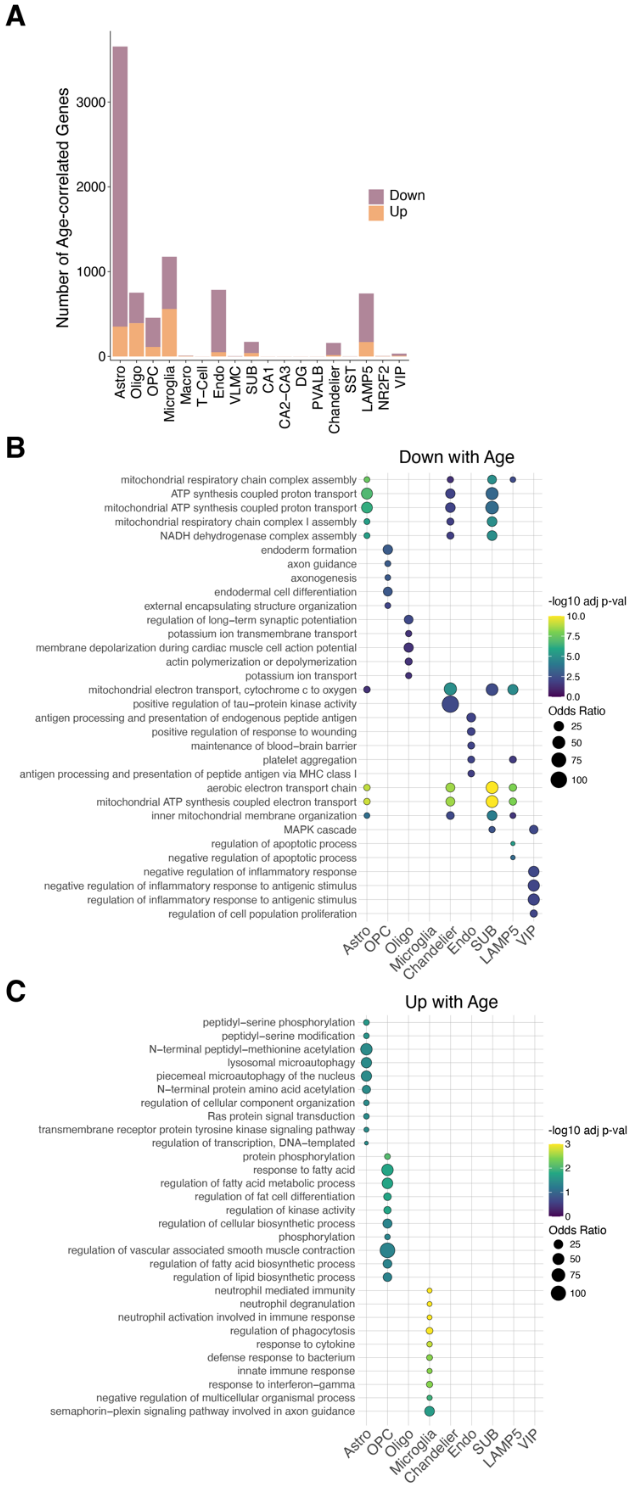


**Fig. S5.**
Age-related changes in gene expression across hippocampal cell types. (**A**) Stacked bar plots showing number of significant (FDR < 0.1) age-correlated genes in each subclass. (**B** and **C**) Gene ontology dot plots displaying significant biological process enriched terms for age-correlated genes that decrease with age (B) or increase with age (C). Only cell types with significant terms in either (B) or (C) are displayed.

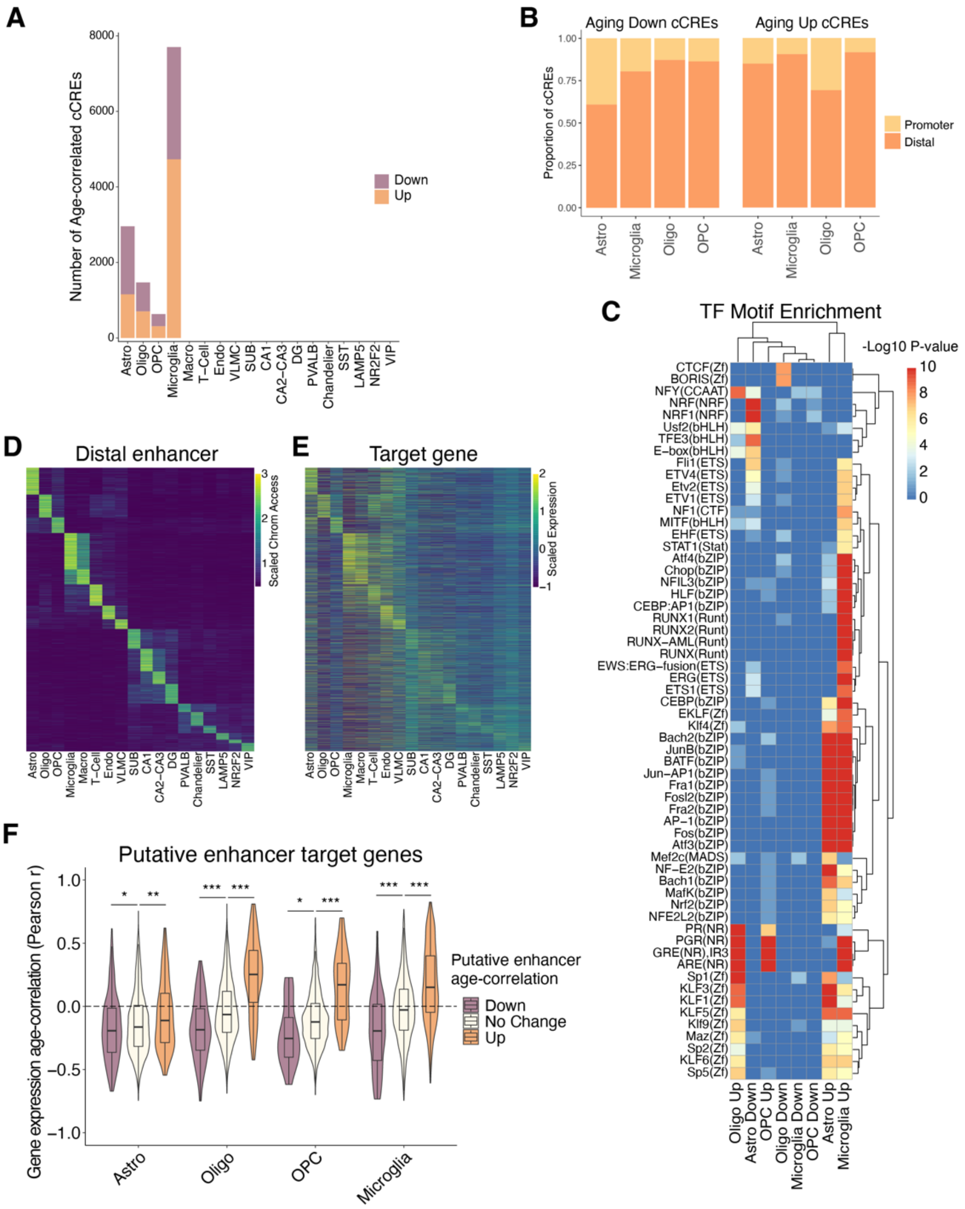


**Fig. S6.**

Aging gene regulatory programs across hippocampal cell types. (**A**) Stacked bar plots showing number of significant (FDR < 0.1) age-correlated cCREs in each subclass. (**B**) Stacked bar plots showing the proportion of age-correlated accessibility of cCREs that decrease (left) or increase (right) with age in promoter and distal genomic regions for four glial cell types (astrocytes, microglia, oligodendrocytes, and OPCs). (**C**) TF motif enrichment heatmap for significant enrichments, q-value < 0.05. "Up" indicates chromatin accessibility increases with age and "Down" indicates chromatin accessibility decreases with age in each subclass. (**D** and **E**) Heatmaps of row-scaled Log2 CPM+1 for chromatin accessibility (D) or gene expression (E) for all cCREs predicted as enhancers by ABC. For each predicted enhancer, the target gene with the highest ABC score is displayed. Rows are clustered by cell type with the highest CPM value of chromatin accessibility. (**F**) Violin plots showing the distribution of Pearson correlation coefficients for age-correlation of expression for target genes of predicted ABC enhancers. P-values from two-sided, unpaired Wilcoxon-rank sum test. * p < 0.05, ** p < 0.01, *** p < 0.001.

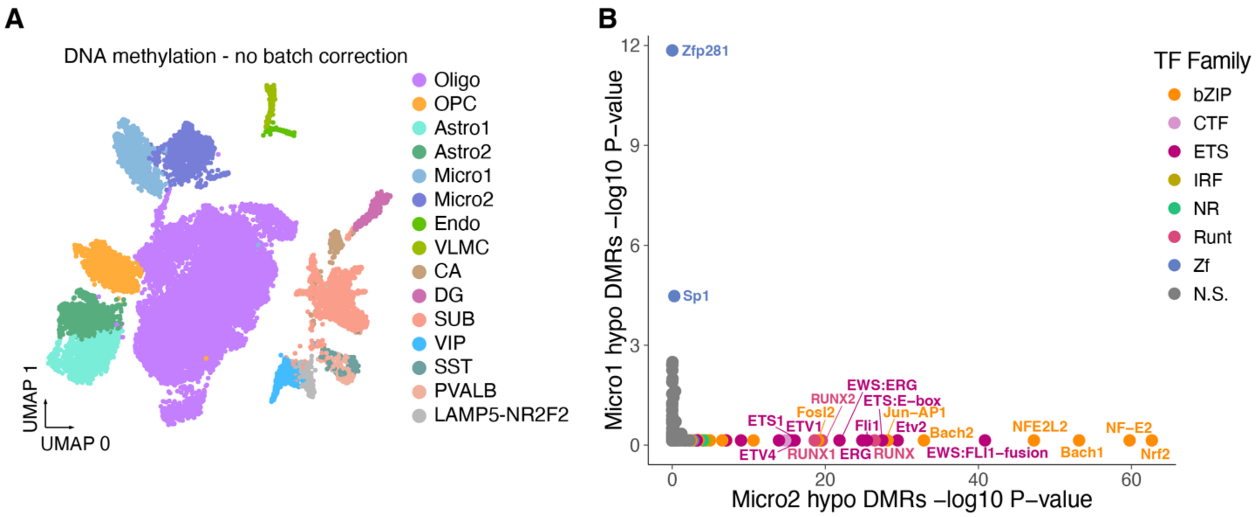


**Fig. S7.**
Microglia subcluster reproducibility and enriched TFs for each subcluster. (**A**) UMAP embedding of snm3C-seq DNA methylation data without batch correction, colored by cell types and subclusters. (**B**) Scatter plot displaying TF motif enrichment in DMRs differentially methylated in either Micro1 or Micro2. Only TF motifs with q-value < 0.05 are colored by its TF family.

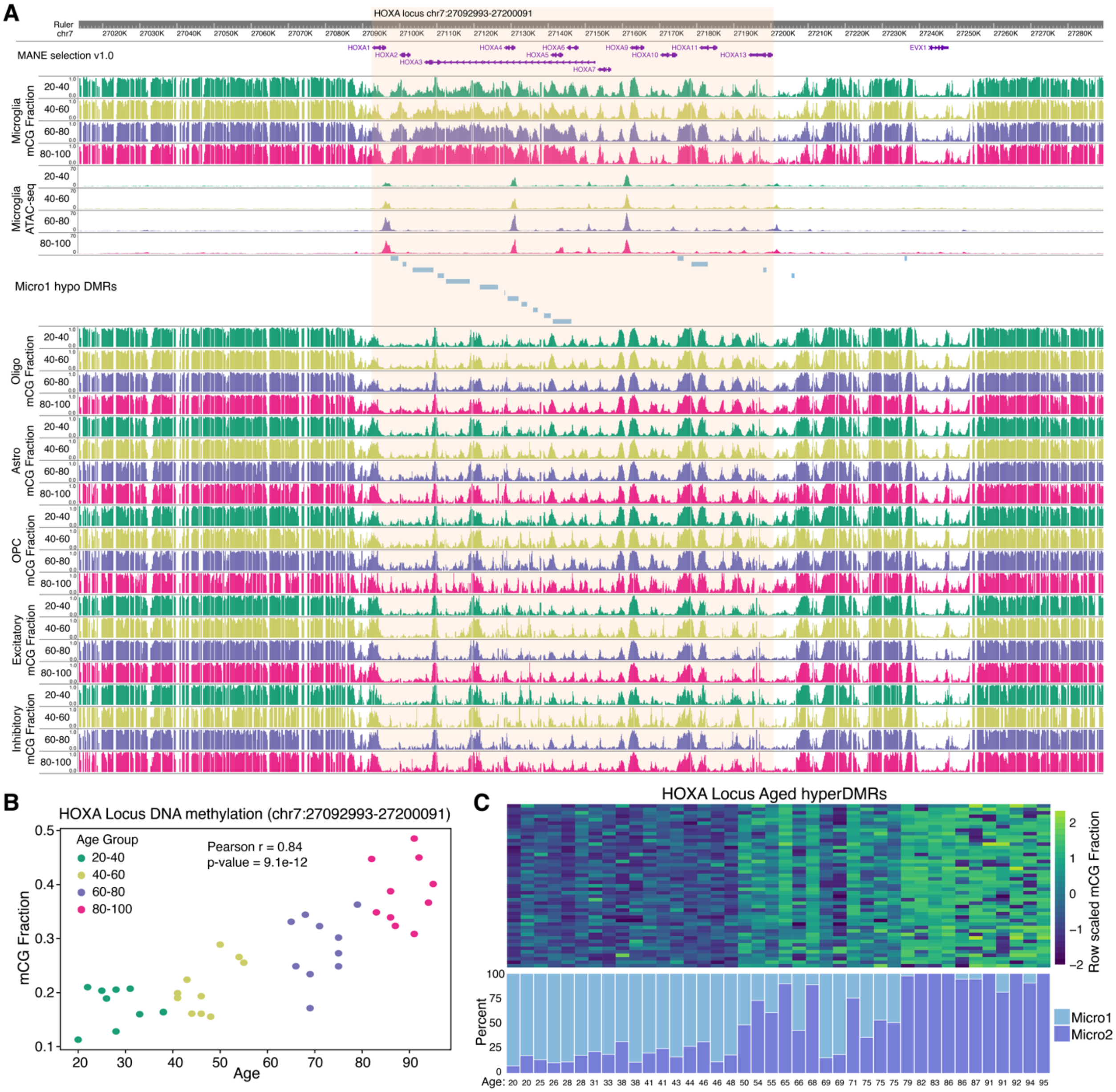


**Fig. S8.**
Characterizing age-correlated DNA methylation in microglia. (**A**) WashU browser snapshots of the HOXA cluster region, showing mCG fraction and chromatin accessibility in microglia across age groups, hypo-methylated regions in Micro1, and mCG fraction in other cell types across age groups. (**B**) Scatter plot showing the mCG fraction of the HOXA locus in microglia from each donor, with each dot representing a donor and colored by age group. (**C**) Heatmap of row-scaled mCG fraction for each donor (columns) with positively age-correlated DNA methylation within the HOXA cluster locus (chr7:27092993-27200091) (top) and stacked bar plots showing the ratio of Micro1 to Micro2 cells recovered for each donor in snm3C-seq experiments (bottom).

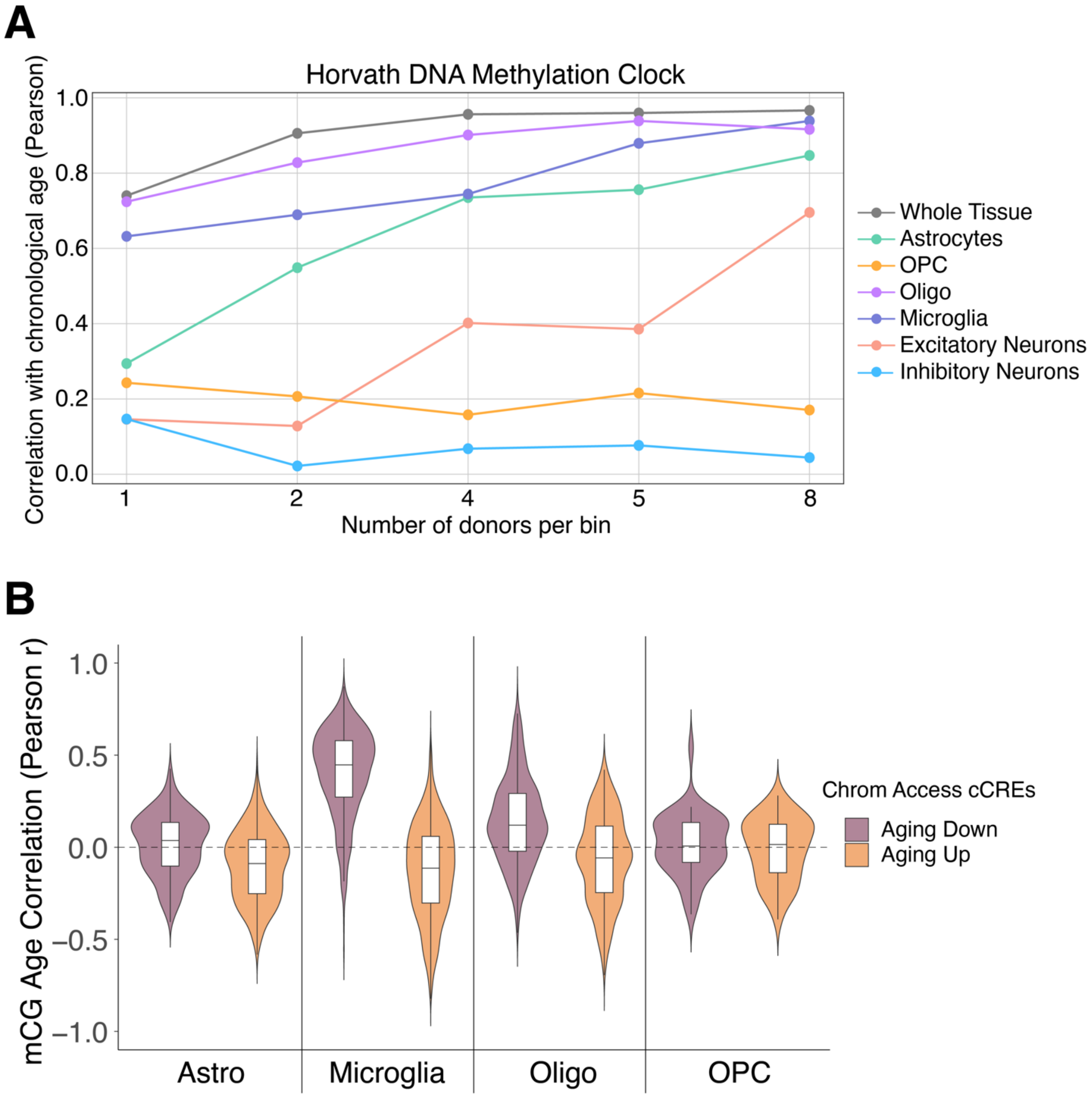


**Fig. S9.**
Aging clock and age-correlated changes in DNA methylation in microglia. (**A**) Pearson correlation coefficients between the Horvath DNA methylation clock (*31*) predicted biological age and chronological age for each subclass with varying numbers of donors per bin. The color of the lines represents different subclasses. The x-axis represents the number of donors per bin, with n=1 indicating correlation across all 40 donors and n=8 representing correlation across 5 age groups using the average age of 8 donors. (**B**) Violin plots showing the distribution of Pearson correlation coefficients for age-correlation of mCG fractions at DMRs overlapping age-correlated cCREs. (**C**) Scatter plot of expression of TEs overlapping Micro2 hypo-methylated marker DMRs vs. donor age for 16 donors as in Fig. 5K.

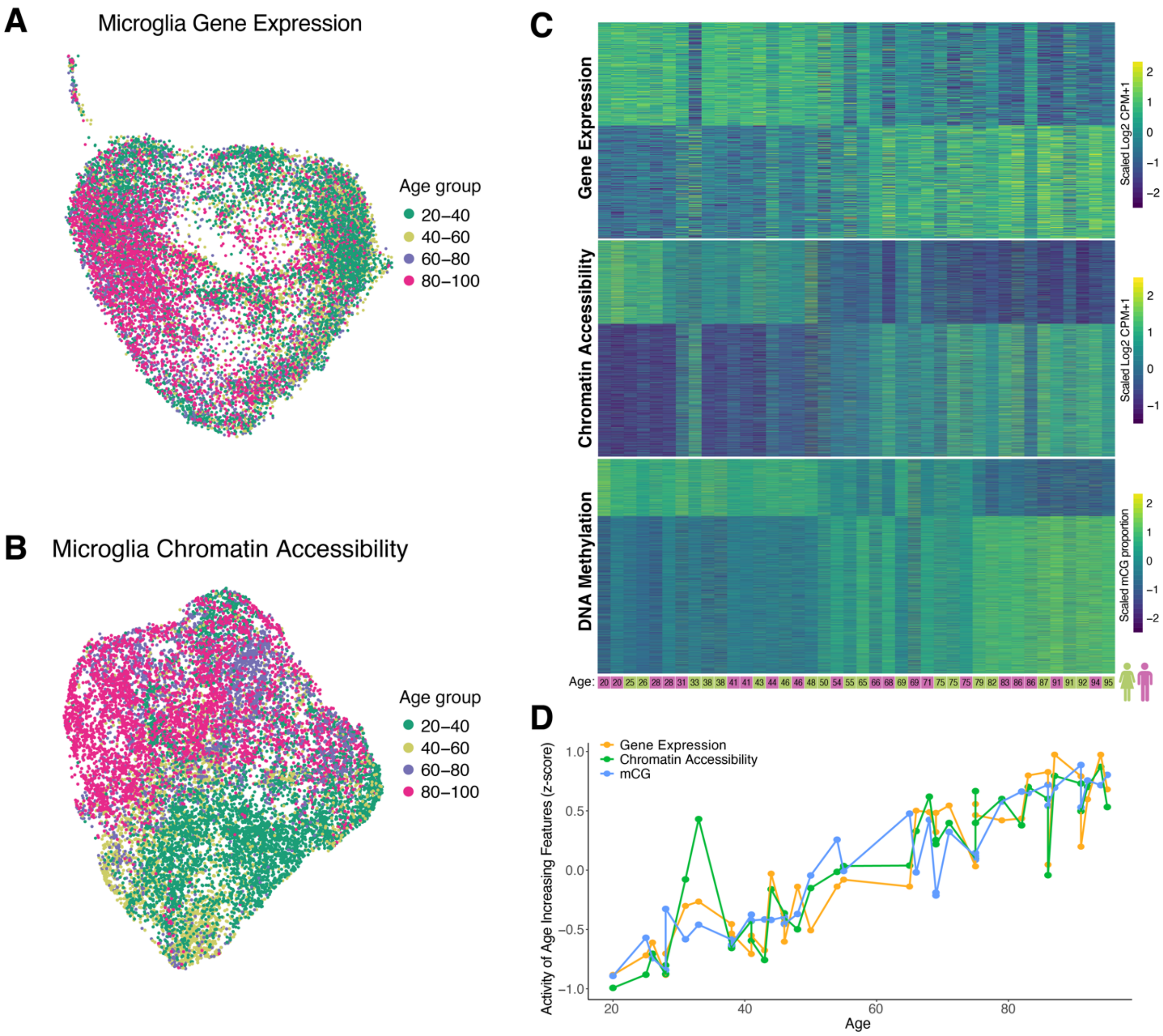


**Fig. S10.**
Age-correlated dynamics of microglia transcriptome, chromatin accessibility, and DNA methylation. (**A**) UMAP displaying microglia 10x multiome clustered by gene expression. Cells are colored by age group. (**B**) Same as (A) except cells are clustered using chromatin accessibility of microglia cCREs. (**C**) Heatmaps displaying row scaled gene expression, chromatin accessibility, and DNA methylation of age-correlated features. Columns represent individual donors, ordered from youngest to oldest with age and sex indicated below. (**D**) Connected scatter plot showing the average values for positively age-correlated features for the indicated data modalities vs age of donor.

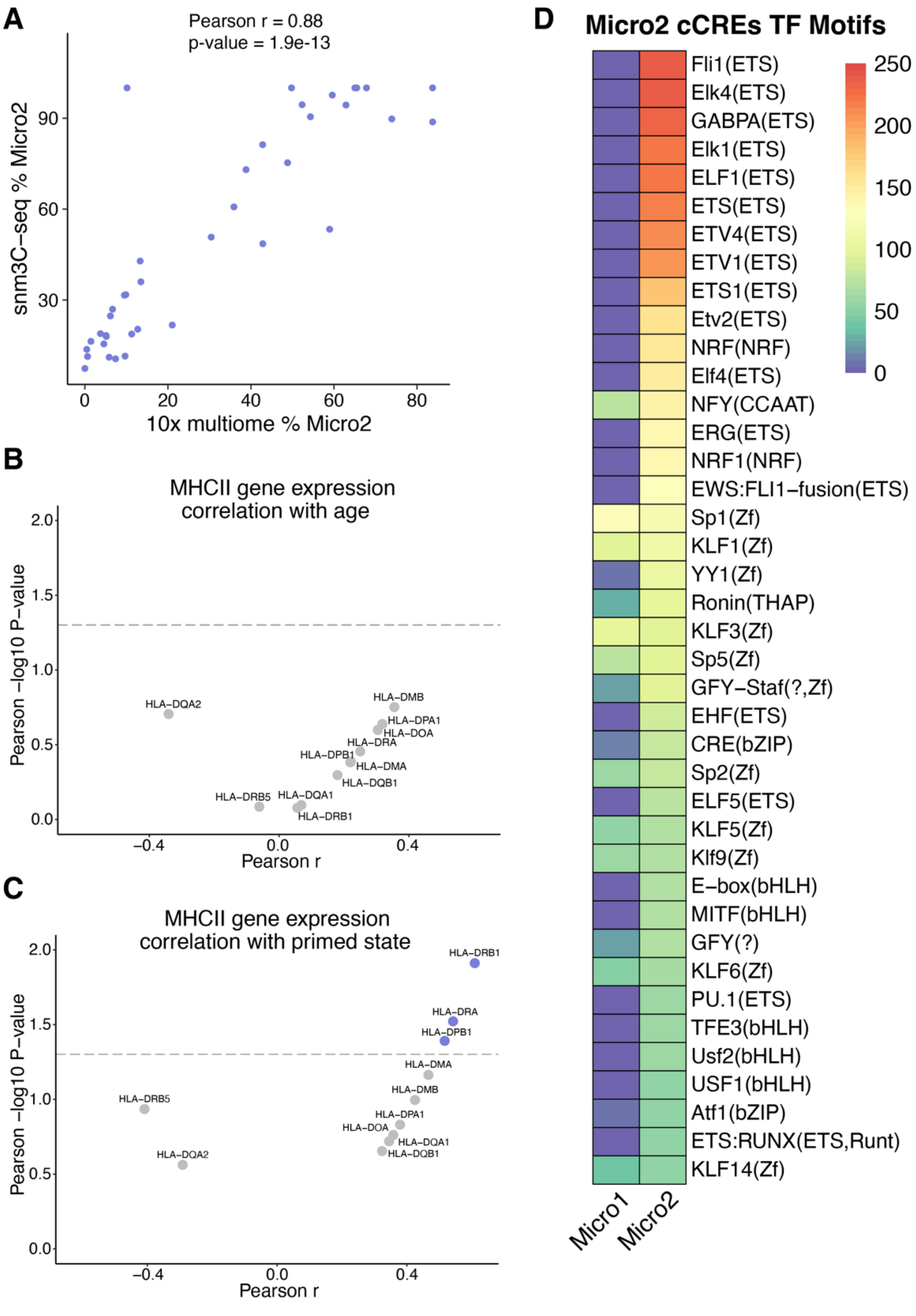


**Fig. S11.**
Identification of primed microglia and characterization of gene expression and chromatin accessibility. (**A**) Scatter plot showing correspondence between percentage of Micro2 (primed microglia) identified in each donor between sequencing technologies (**B**) Scatter plot showing correlation of expression of MHC II genes with age or (**C**) percentage of Micro2 cells (primed state) captured from snm3C-seq assay for each donor. (**D**) TF motif enrichment heatmap for significant enrichments of differential cCREs between Micro1 and Micro2, displaying top 40 TFs ordered by smallest q-value in Micro2 cluster.

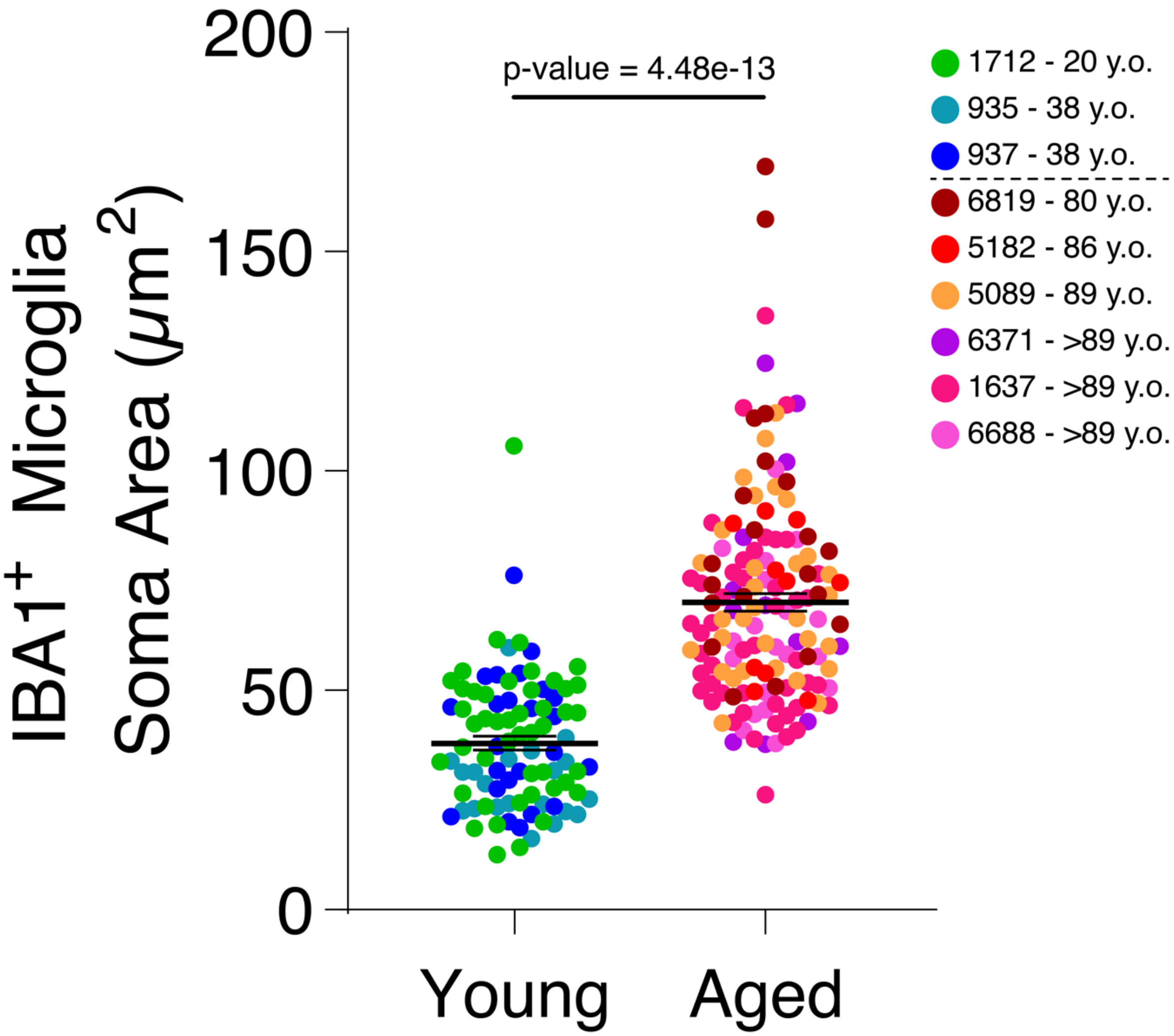


**Fig. S12.**
Aged microglia soma size quantification. Dot plot showing soma size quantification from CA3 region of young (n=3; 20-38 y.o.) and aged (n=6; 80-106 y.o.). Dots represent individual cells (young n=90, aged n=140) and colored by donor. P-value from linear mixed-effect model (LME).

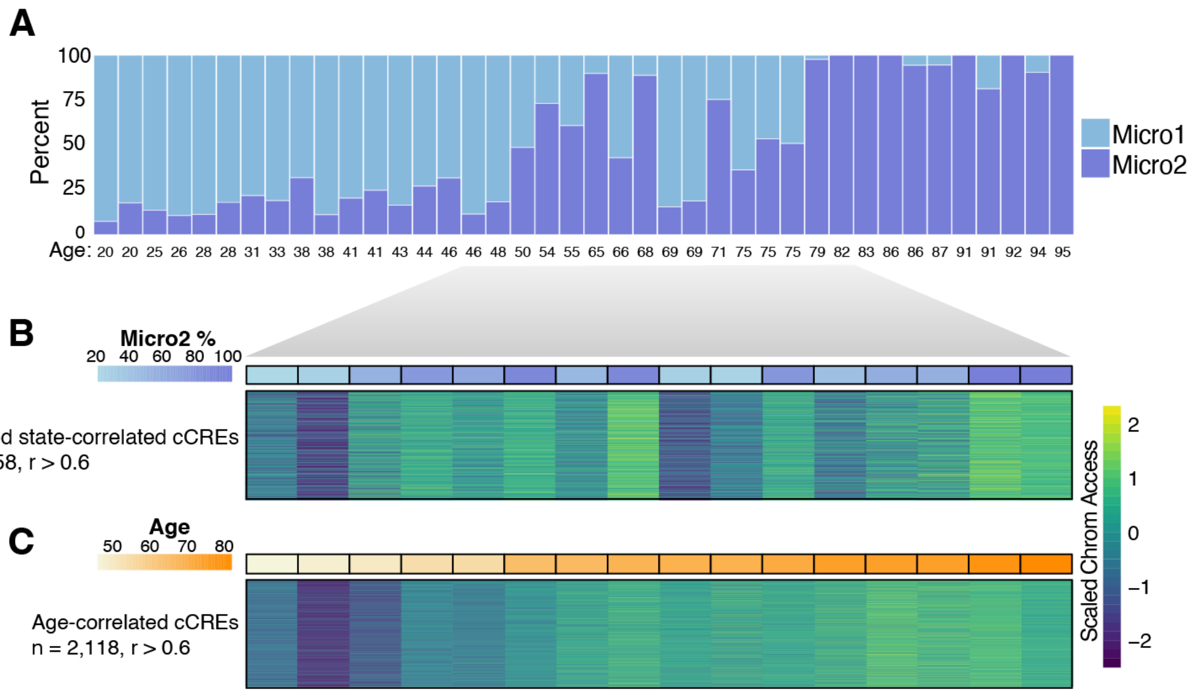


**Fig. S13.**
Microglia primed state and age-correlated cCREs. (**A**) Stacked bar plots showing proportion of Micro1 and Micro2 cells recovered in each donor of snm3C-seq datasets, ordered from youngest to oldest. (**B** and **C**) For 16 donors, ages 46 to 82 where age does not correlate with microglia subcluster proportions, heatmap displays their microglia chromatin accessibility (scaled Log2 CPM+1) for cCREs that correlate with Micro2 percentage across these 16 donors ("Primed state-correlated cCREs"), PCC > 0.6 (B) and heatmap displaying the chromatin accessibility (scaled Log2 CPM+1) for cCREs that correlate with age across these 16 donors, PCC > 0.6 (C).

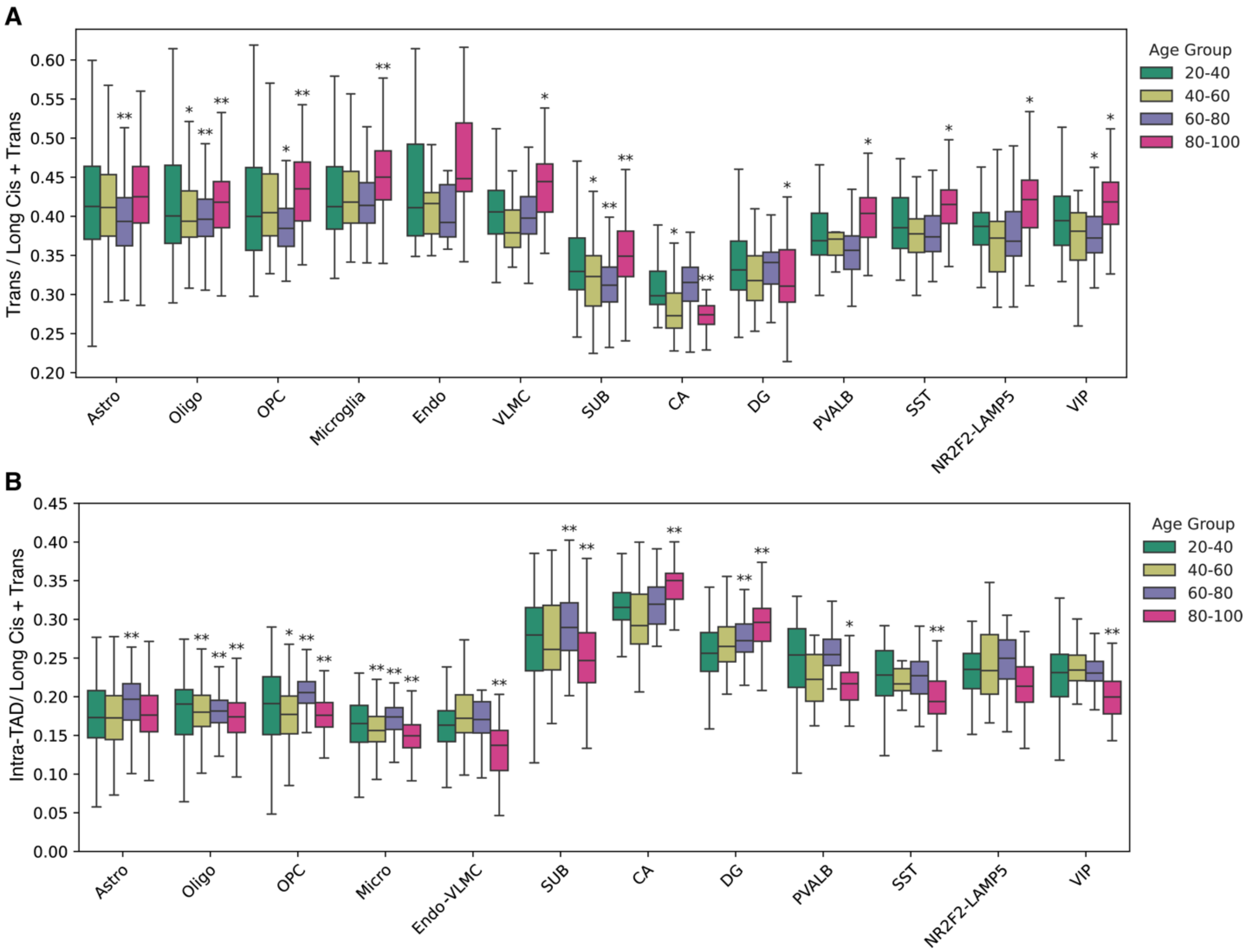


**Fig. S14.**
Restructuring of the aging 3D genome. (**A**) Box plots displaying the ratio of trans contacts to cis long and trans contacts for cell types across age groups. (**B**) Box plots displaying the ratio of intra-TAD contacts to cis long and trans contacts for cell types across age groups.

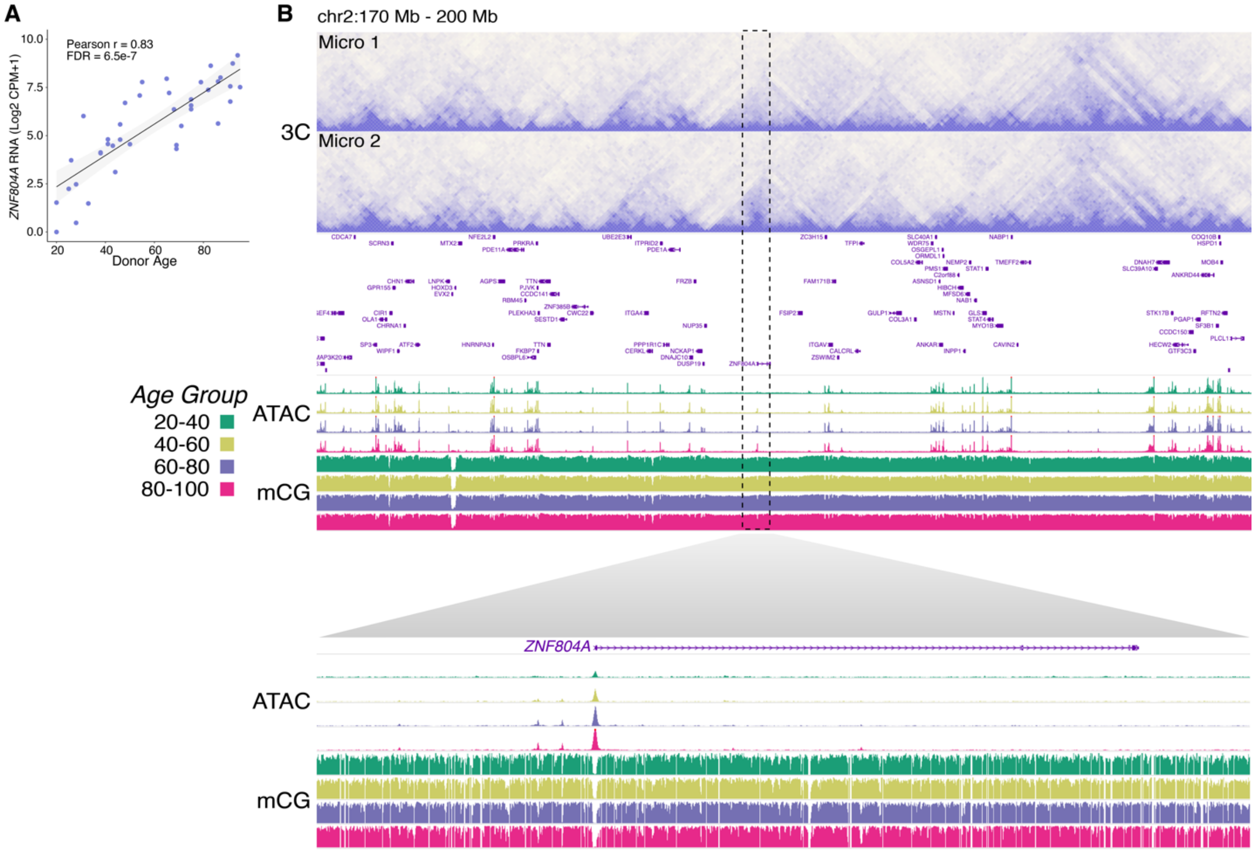


**Fig. S15.**
Age-correlated expression and chromatin restructuring at *ZNF804A*. WashU Epigenome browser snapshots of the chr2:170Mb-200Mb region. The heatmap on the top shows contact frequency in Micro1 and Micro2. Below the heatmap, plots display chromatin accessibility and mCG fraction in microglia across age groups. A dotted box highlights the *ZNF804A* region, with a zoomed-in view of chromatin accessibility and mCG fraction in this region across age groups provided below.

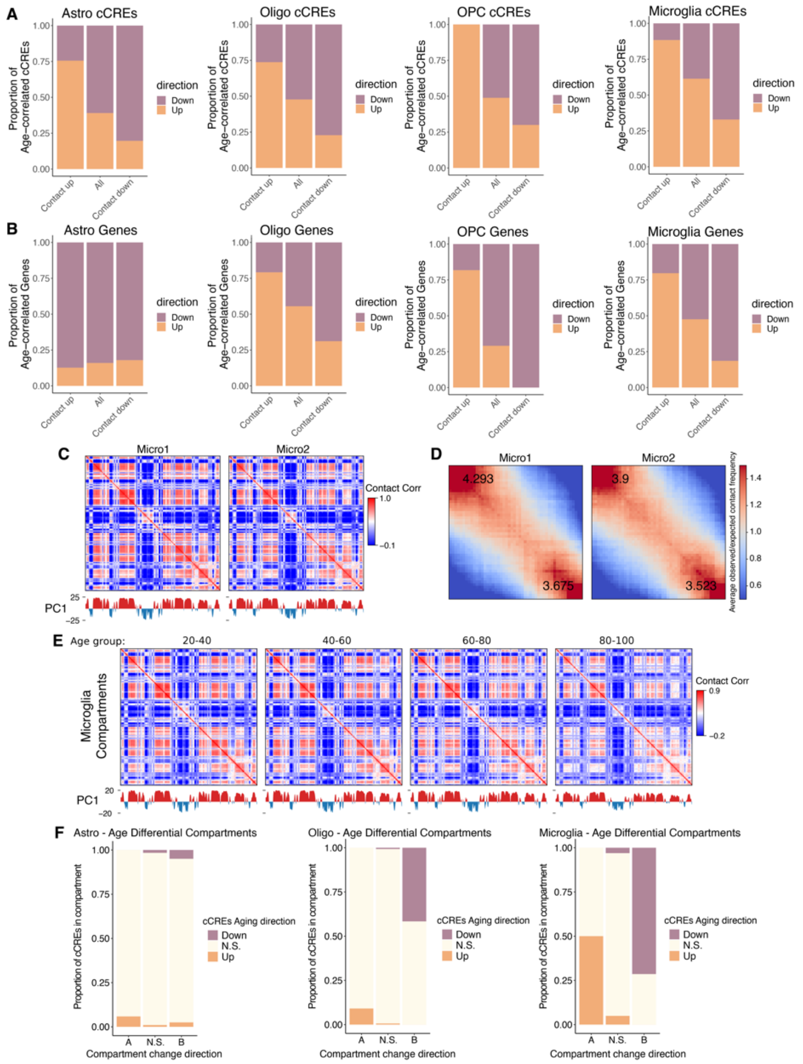


**Fig. S16.**
Correspondence between dynamic 3D genome architecture, the epigenome, and transcriptome during aging. (**A** and **B**) Stacked bar plots show proportions of age-correlated cCREs (A) and age-correlated genes (B) within differential contact bins for each subclass. (**C**) Heatmaps displaying the correlation matrices of distance-normalized contact maps for Micro1 and Micro2 at a 100kb resolution for the region chr2:150Mb-200Mb. Line plots representing PC1 of these correlation matrices (bottom). (**D**) Saddle plots showing average observed over expected contact frequency between inactive compartments (BB interactions; in the upper left), active compartments (AA interactions; in the lower right), and active compartments and inactive compartments (AB and BA; in the bottom left and top right). The values represent the strength calculated as the ratio of AA interactions to AB interactions and the ratio of BB interactions to BA interactions. (**E**) Heatmaps showing the correlation matches of distance-normalized contact maps for each age group in microglia, after downsampling to the lowest number of total contacts among the age groups, at a 100kb resolution for the region chr2:150Mb-200Mb. Line plots displaying PC1 of these correlation matrices (bottom). (**F**) Stacked bar plots show proportions of age-correlated cCREs or not significantly age-correlated (N.S.) in differential compartments that became more active ('A' on x-axis'), or more inactive ('B' on x-axis), or not significantly differential ('N.S.' on x-axis).

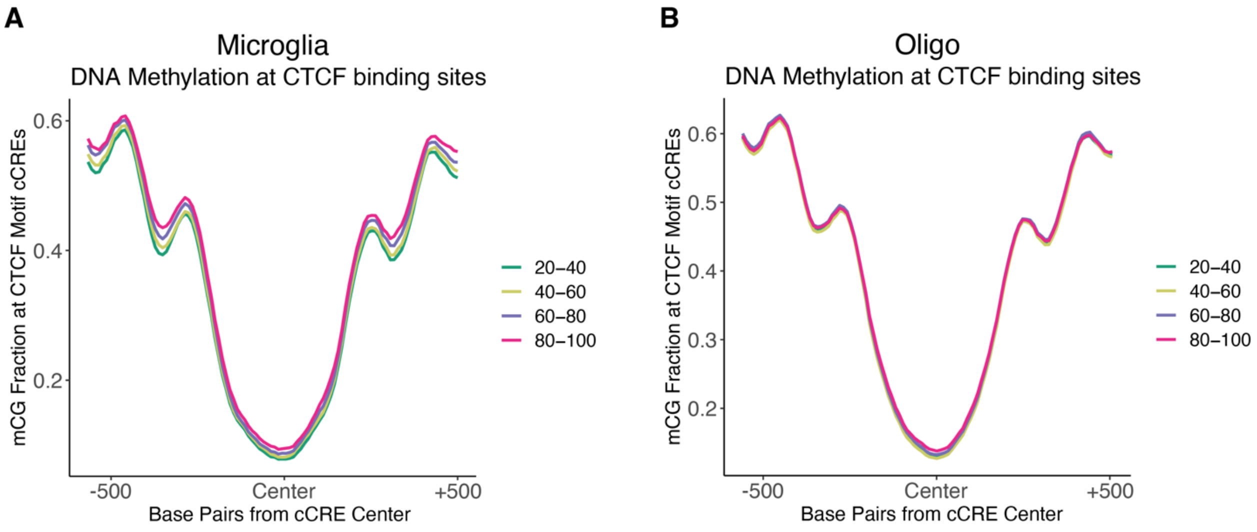


**Fig. S17.**
DNA methylation profiles at CTCF motifs with age. (**A**) Average mCG fractions at CTCF sites in each age group in microglia and (**B**) oligodendrocytes.

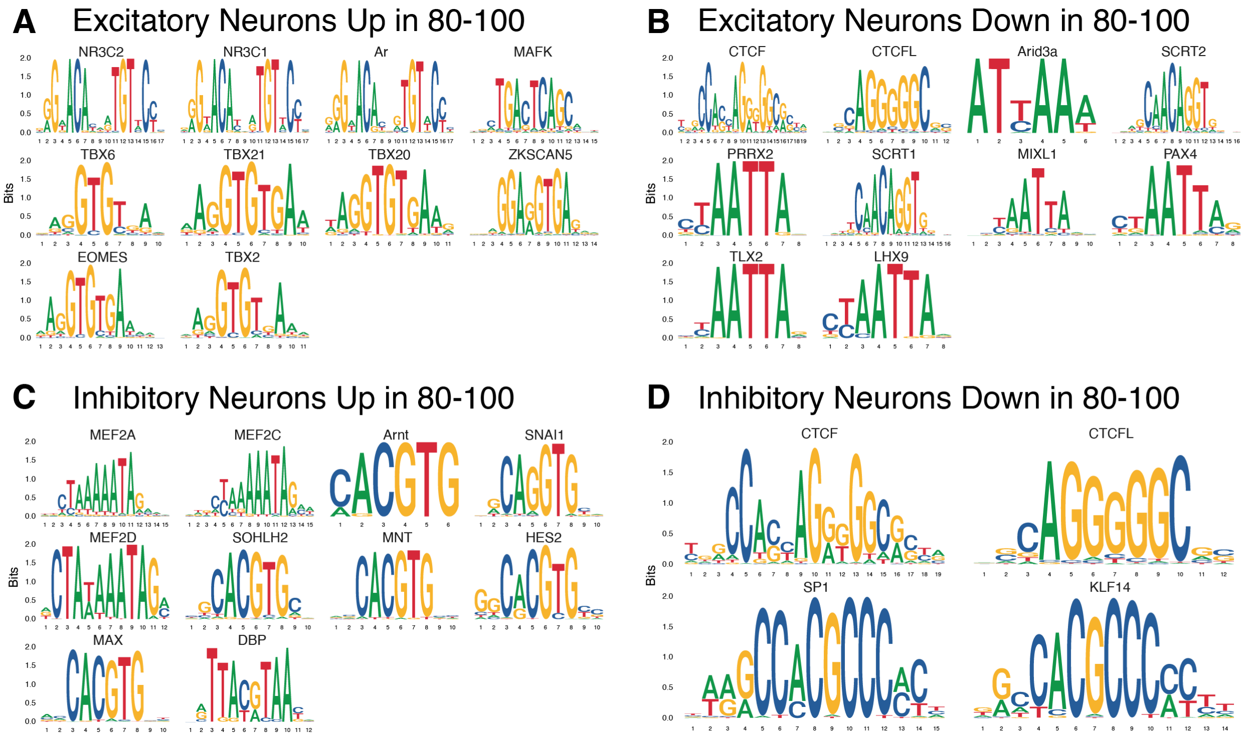


**Fig. S18.**
Age-correlated TF motif enrichment in neurons. (**A** to **D**) TF motif enrichment at differentially enriched ATAC-seq fragments between 80-100 year old and 20-40 year old excitatory or inhibitory neurons.
